# Hierarchical reactivation of transcription during mitosis-to-G1 transition by Brn2 and Ascl1 in neural stem cells

**DOI:** 10.1101/2021.02.17.431407

**Authors:** Mário A. F. Soares, Diogo S. Soares, Vera Teixeira, Raul Bardini Bressan, Steven M. Pollard, Raquel A. Oliveira, Diogo S. Castro

## Abstract

During mitosis, chromatin condensation is accompanied by a global arrest of transcription. Recent studies suggest transcriptional reactivation upon mitotic exit occurs in temporally coordinated waves, but the underlying regulatory principles have yet to be elucidated. In particular, the contribution of sequence-specific transcription factors (TFs) remains poorly understood. Here we report that Brn2, an important regulator of neural stem cell identity, associates with condensed chromatin throughout cell division, as assessed by live-cell imaging of proliferating neural stem cells. By contrast, the neuronal fate determinant Ascl1 dissociates from mitotic chromosomes. ChIP-seq analysis reveals that Brn2 mitotic-chromosome binding does not result in sequence-specific interactions prior to mitotic exit, relying mostly on electrostatic forces. Nevertheless, surveying active transcription using single-molecule RNA-FISH against immature transcripts, indicates the differential presence of TF near chromatin when exiting mitosis is associated with early (anaphase) versus late (early G1) reactivation of key targets of Brn2 and Ascl1, respectively. Moreover, by using a mitotic-specific dominant negative approach, we show that competing with Brn2 binding during mitotic exit reduces the transcription of its target gene Nestin. Our study shows an important role for differential binding of TFs to mitotic chromosomes, governed by their electrostatic properties, in defining the temporal order of transcriptional reactivation during mitosis-to-G1 transition.

## Introduction

During cell division complex cellular changes occur that have a dramatic impact on genome regulation. This is particularly the case in mitosis, as the core transcriptional machinery is heavily inhibited, and transcription globally arrested (Steffen and Ringrose 2014; Palozola, Lerner, and Zaret 2019). Condensation of chromatin and the dilution of nuclear proteins occurs following nuclear envelope break-down (NEBD), altering the activities of DNA-binding transcription factors (TFs). Nevertheless, in recent years several TFs have been reported to occupy a fraction of their genomic target sites during mitosis (Kadauke et al. 2012; Caravaca et al. 2013; Festuccia et al. 2016; Deluz et al. 2016; Liu et al. 2017). This process, referred to as “mitotic bookmarking”, provides an attractive model for how gene regulatory information is propagated across cell divisions. TF binding to regulatory regions may therefore mark a subset of genes for efficient reactivation upon mitotic exit (Festuccia et al. 2017; Raccaud and Suter 2018; Palozola, Lerner, and Zaret 2019). One possibility is that occupancy of regulatory regions by TFs maintains nucleosome positioning, counteracting large chromatin rearrangements characteristic of mitosis, as shown in recent studies (Festuccia et al. 2019; Owens et al. 2019). Nevertheless, the basis and impact of mitotic chromosome binding by TFs to gene regulatory events remains poorly understood.

Distinct types of interactions mediate the association of TFs with DNA. Long-lasting sequence-specific interactions occur via direct binding to specific nucleotide motifs, and can be mapped using chromatin immunoprecipitation followed by sequencing (ChIP-seq). In addition, TFs take part in non-specific interactions with chromatin, mediated primarily by electrostatic forces that do not depend on specific DNA sequences (Suter 2020). These result from the strong electrostatic field surrounding negatively charged DNA (even when in a nucleosome), and are of most importance when chromatin is highly compacted, as it is the case of condensed chromosomes during mitosis, or heterochromatic regions in the interphase nucleus (Festuccia et al. 2019; Raccaud et al. 2019; Gebala et al. 2019). Mitotic bookmarking implies site specific binding (i.e. direct interactions with the specific DNA motifs); however, non-specific interactions could also contribute to this process (Caravaca et al. 2013). In addition, because many TFs that bind mitotic chromatin are also pioneer TFs (i.e. bind to nucleosomal DNA), the two activities have been often associated in the literature (Zaret 2014).

Recent studies, performed in a limited number of cell types, suggest reactivation of transcription upon mitotic exit does not occur in bulk during G1, but in sequential waves with distinct kinetics during mitosis-to-G1 (M-G1) transition (Palozola et al. 2017; Abramo et al. 2019; Zhang et al. 2019; Kang et al. 2020). How are such distinct temporal profiles of gene reactivation established? Some studies have suggested sequence-specific binding of TFs to mitotic chromatin as the basis for early reactivation of bookmarked genes during M-G1 (Kadauke et al. 2012; Caravaca et al. 2013; Festuccia et al. 2016; Liu et al. 2017). However, these observations are still controversial, due to the lack of temporal resolution and sensitivity of cell-population assays used to survey active transcription during M-G1. In addition, they also fail to compare TFs with distinct mitotic-binding abilities.

Given the view that mitosis (and in particular the M-G1 transition) is a window of opportunity for cell state transitions, recent reports on the interplay between TFs and mitotic chromatin have been performed in the context of pluripotent Embryonic Stem (ES) cells, which have unusual cell cycle and chromatin features compared to somatic cells. By contrast, no studies have so far investigated mitotic TF binding in neural stem cells, where in vitro models and knowledge of the key cell fate determinants are available.

Members of the Class III POU family of TFs (e.g. Brn2 - aka Pou3f2) play diverse functions in neural development, including the acquisition and maintenance of neural progenitor identity. They function by directly binding and regulating a large number of neural stem cell enhancers, often in partnership with Sox2, another TF critical for maintenance of neural stem/progenitor cells (Tanaka et al. 2004; Miyagi et al. 2006; Lodato et al. 2013). Commitment towards the neuronal fate is primarily regulated by basic-helix-loop-helix (bHLH) TFs of the proneural family such as Ascl1 (also termed Mash1), which is both necessary and sufficient to activate a full program of neuronal differentiation (Bertrand, Castro, and Guillemot 2002; Vasconcelos and Castro 2014). Such function during development is reinforced by the extensive use of Ascl1 in reprogramming somatic cells into induced-neurons, which has been attributed to its pioneer TF activity. In addition to a pivotal role in triggering neuronal differentiation, Ascl1 is also required for proper proliferation of Radial Glia (RG) neural stem cells in the embryonic brain, and in RG-like cultures growing in non-differentiating conditions (Castro et al. 2011; Imayoshi et al. 2013). Activation of neuronal differentiation by proneural factors requires (and occurs concomitantly with) the suppression of a progenitor program, of which Sox2 and Brn2 are major regulators (Bylund et al. 2003; Vasconcelos et al. 2016). The balance between these opposing TFs is therefore critical in determining, whether upon cell division, newborn cells maintain their progenitor identity or instead become committed to neuronal differentiation.

Here, we explore the relevance of mitotic bookmarking mechanisms by TFs in neural stem/progenitor cells, focusing on Brn2 and Ascl1, given their critical role in neural stem cell maintenance and differentiation, as well as in neural lineage specific expression. By combining live-cell imaging, genome-wide location analysis by ChIP-seq and single-molecule RNA fluorescence *in situ* hybridization (smRNA-FISH), we addressed how these two pivotal TFs interact with mitotic chromatin, and how this impacts their gene regulation functions upon mitotic exit.

## Results

### Brn2 and Ascl1 display distinct mitotic chromosome binding abilities

Live-cell imaging microscopy has recently emerged as an important method for probing the association of TFs with condensed chromosomes, circumventing artifacts caused by chemical fixation (Teves et al. 2016). We applied this imaging method to cultured RG-like neural stem/progenitor cells originated from embryonic mouse telencephalon (herein referred to as NS cells), undergoing proliferative divisions under non-differentiating conditions. Tagging of endogenous Brn2 and Ascl1 with enhanced green fluorescent protein (eGFP) allowed surveying their dynamics of association with chromatin, without altering expression levels (Figure 1A-C, E). Strikingly, imaging of Brn2 in NS cells reveals this TF associates with mitotic chromosomes during all stages of mitosis, as shown by strong colocalization of Brn2-eGFP with DNA (Figure 1B). The degree of colocalization during metaphase was quantified by determining the level of mitotic chromosome enrichment (MCE; metaphase plate fluorescence divided by total fluorescence level) (Figure 1D). By contrast, the same approach used with Ascl1 reveals clear exclusion of eGFP signal from condensed chromatin since NEBD (i.e. prometaphase), with eGFP signal dispersing within the mixed nucleoplasm and cytoplasm (Figure 1C). Exclusion from chromatin persisted until cytokinesis (telophase: metaphase + 10 min), with colocalization of eGFP with DNA becoming apparent from late telophase onwards (metaphase + 15 min) (Figure 1C-D).

**Figure 1.**
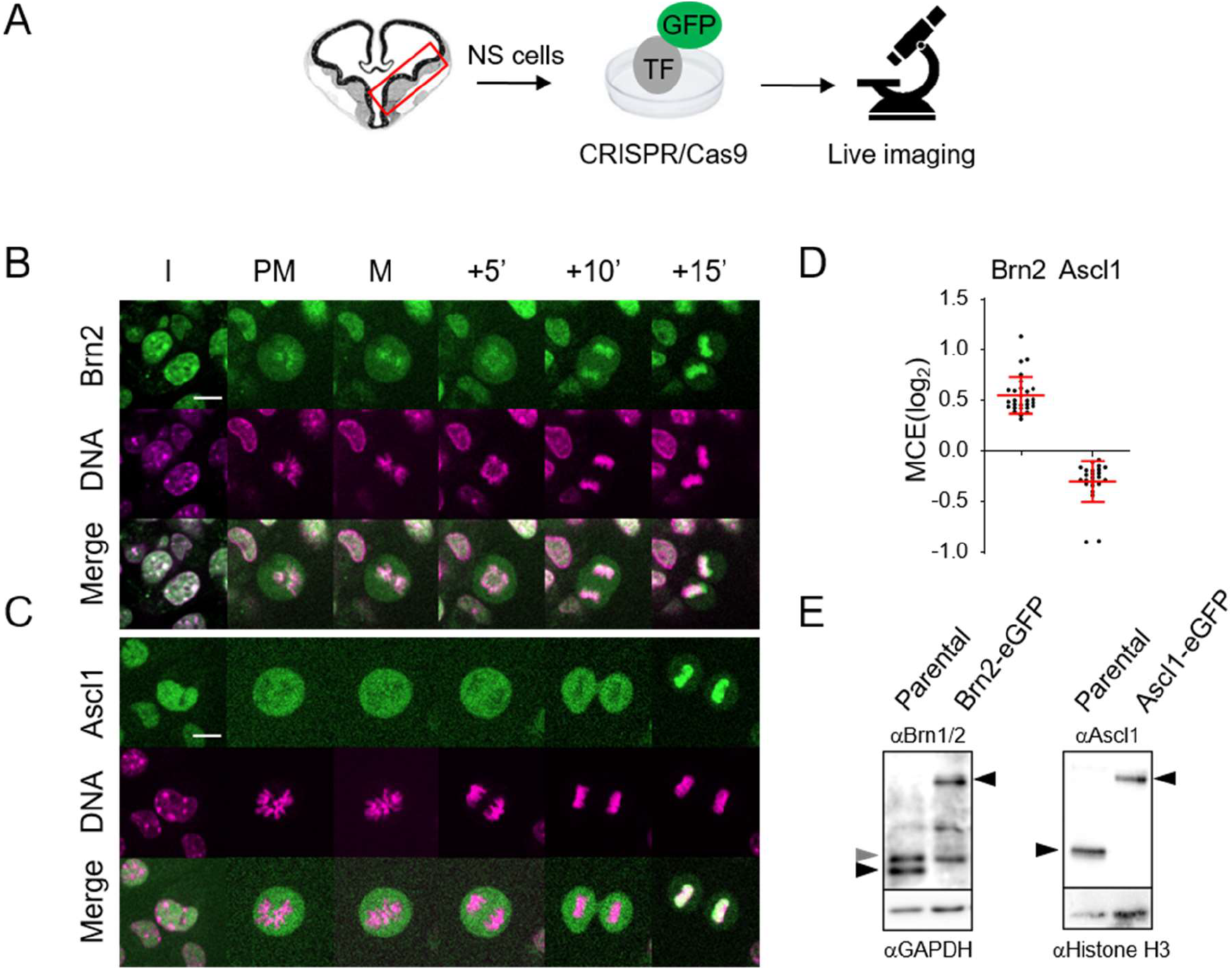
Brn2 and Ascl1 display distinct interactions with condensed chromosomes in proliferating NS cells. **(A)** Adherent cultures of mouse embryonic neural stem/progenitors of ventral telencephalon expressing eGFP tagged Ascl1 or Brn2 were live imaged undergoing proliferative divisions. **(B-C)** Time-lapse live-cell imaging of NS cells expressing eGFP tagged Ascl1 (A) and Brn2 (B) in presence of DNA dye SiR-Hoechst. **(D)** Quantifications of mitotic chromosome enrichment levels during metaphase in NS cells expressing Ascl1-eGFP (n=24) and Brn2-eGFP (n=28). Data shown as mean ± SD. **(E)** Western-blot analysis of Ascl1-eGFP and Brn2-eGFP protein expression in lines used in live imaging, and their counterparts in parental NS cells (See Supplemental Figure S1 for characterization of Brn2 commercial antibody). Black arrows mark specific bands corresponding to endogenous or eGFP fusion proteins. Grey arrow marks Brn1, also recognized by the antibody used (see Supplemental Figure S2). Scale bar=10μm.

Colocalization of Ascl1 with DNA in late telophase may be indicative of binding to chromatin, or instead result from active import of Ascl1-eGFP into newly formed nuclear envelope, wrapping tightly around decondensing chromosomes. To address this, we started by mapping the Ascl1 nuclear localization signal (NLS). A 23 residues long sequence N-terminal from the bHLH domain, highly predicted as a bipartite NLS, is both required (when mutated in context of Ascl1-eGFP), and sufficient (when fused to eGFP), to drive nuclear localization (Supplemental Figure S2). Importantly, the timing of association of Ascl1-eGFP with chromatin in dividing P19 embryonic carcinoma cells, where the analysis of these Ascl1 derivatives was performed, parallels that observed in NS cells. Strikingly, mutating the Ascl1 NLS in one part, or both parts of the NLS sequence, results in a significant delay (>60 min) or complete inability to colocalize with DNA at mitotic exit, respectively (Supplemental Figure S2C). In conclusion, Brn2 and Ascl1 have contrasting binding patterns through mitosis. Colocalization of Ascl1 with DNA in late telophase results from its active import into the nuclear envelope rather than its ability to bind to mitotic chromatin prior to nuclear envelope reformation. The observed binding of Brn2 may suggest some role as mitotic bookmarker.

### Ascl1 protein is present during M-G1 transition in vivo

TFs can be targeted for degradation during mitosis, imposing another layer of regulation of their potential eviction from mitotic chromatin (Festuccia et al. 2017). We next ascertained if Ascl1 protein can also be detected during mitosis in an *in vivo* context. The expression of endogenous

Ascl1 during mitosis was characterized by immunohistochemistry in the developing ventral telencephalon at E12.5, a developmental stage when most mitotic events occur in apically dividing progenitors, as seen by phospho-histone H3 staining (Figure 2A). Most of these are RG that divide asymmetrically to self-renew and originate another more differentiated progenitor (e.g. short neural precursor) (Pilz et al. 2013). Dynamic expression of Ascl1 driven by Hes1 oscillations (Imayoshi et al. 2013) results in heterogeneous expression in neural stem/progenitor cells in both germinal layers (ventricular and subventricular zones – VZ and SVZ) (Figure 2B). Ascl1 protein is detectable in 64.9% of apical mitotic cells, which can be segregated by their high or low Ascl1 protein levels. Using DAPI to assess cell-cycle phase, it is observed that Ascl1 expression (in both high and low expressing cells) remains constant until anaphase, with fewer (but still significant number of cells) in telophase retaining Ascl1 expression (34.9%) (Figure 2B-D). Mitotic cells expressing moderate to high levels of Ascl1 were also observed in other VZ (sub-apically dividing) and SVZ progenitors undergoing mitosis (data not shown). Altogether, these observations suggest that a significant number of neural progenitor cells maintain moderate levels of Ascl1 throughout M-phase, although Ascl1 protein levels are reduced upon mitotic exit. We therefore conclude that protein degradation is not the key pathway that limits Ascl1 binding to mitotic chromatin.

**Figure 2.**
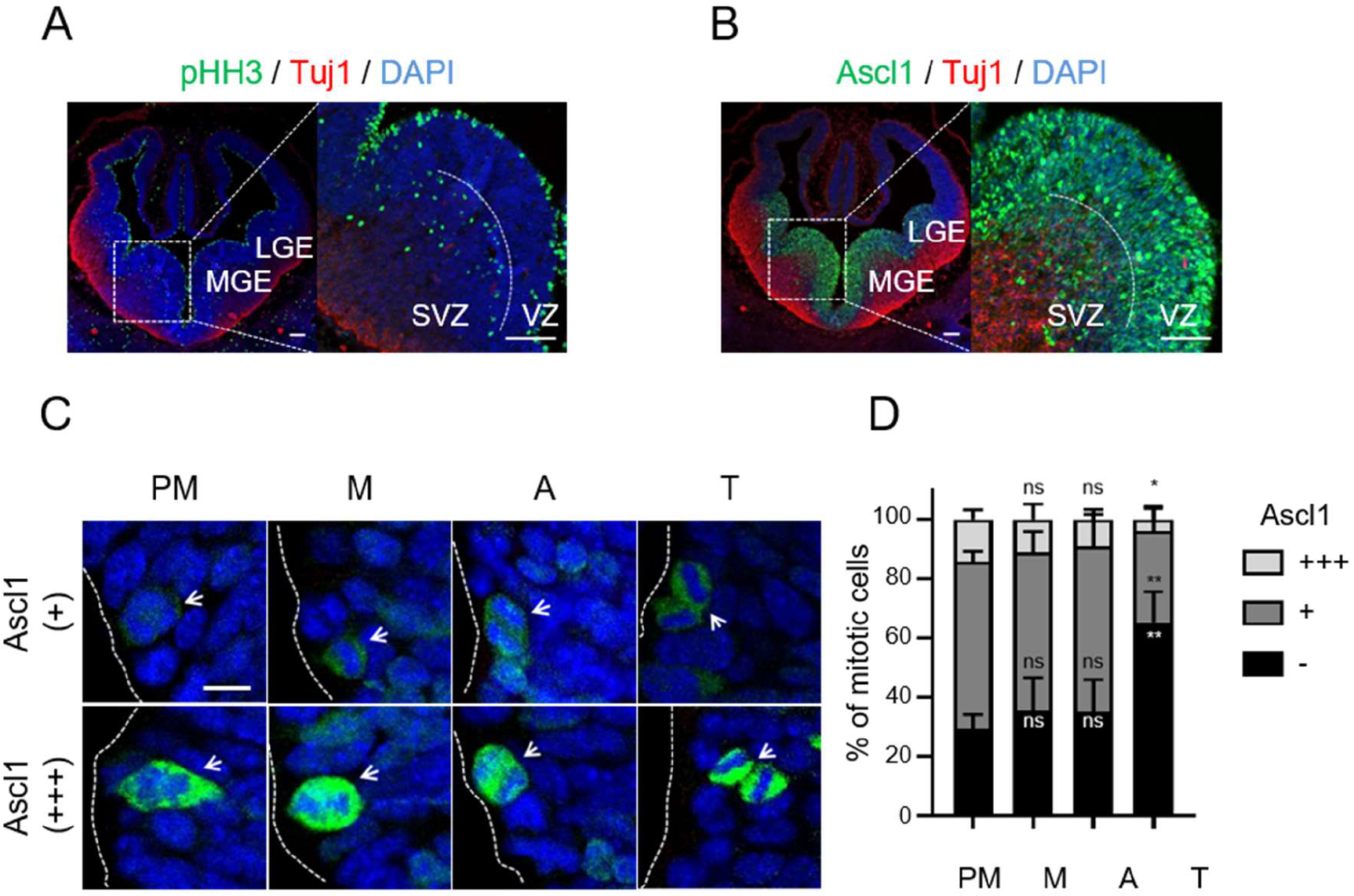
Quantification of Ascl1 expression in apically dividing neural progenitors of ventral telencephalon. **(A-B)** Cross-sections of mouse telencephalon immuno-stained for pHH3 and Tuj1 (A) or Ascl1 and Tuj1 (B) together with DAPI staining. Mitotic pHH3^+^ cells can be found in the ventricular zone (VZ) and subventricular zone (SVZ) in both the lateral and medial ganglionic eminences (LGE and MGE, respectively) of E12.5 ventral telencephalon. **(C)** The cell cycle stage of apically dividing progenitors, both from LGE and MGE, was assessed using DAPI staining, and segregated in cells not expressing (−), expressing low levels (+) or expressing high levels (+++) of Ascl1 protein. **(D)** Stacked bar plots showing the percentage of mitotic cells expressing different levels of Ascl1 in prometaphase (PM), metaphase (M), anaphase (A) and telophase (T). Data shown as mean percentage ± SD. Data information: Cross-sections from 3 mice were used. From each mouse, 6 consecutive slices were acquired in a total of 1736 cells being quantified. One-way ANOVA Tukey’s multiple comparison was used to compare Ascl1 levels between prometaphase and other M phase stages. p>0.05 (n.s.), p≤0.05 (*), p≤0.01 (**), p≤0.001 (***), p≤0.0001 (****). Scale bars = 100 μm (A-B) and 5 μm (C).

### eGFP does not hinder Ascl1 mitotic chromosome binding

Ascl1 binds DNA as a heterodimer with ubiquitously expressed bHLH E-proteins (e.g. E47). One possibility could be that the absence of its heterodimeric partner during mitosis hampers Ascl1 binding. However, co-transfecting Ascl1 with E47, or tethering of both proteins via a glycine bridge (Ascl1/E47) (Geoffroy et al. 2009), did not promote its association with mitotic chromosomes (data not shown and Figure 3A and D). To exclude the possibility that the bulkiness of eGFP (~27kDa) hinders the ability of Ascl1 (~30kDa) to interact with mitotic chromatin, we performed live-cell imaging analysis of Ascl1/E47 tagged with a small (12-residues long) tetracysteine tag (TC-Ascl1/E47) (Martin et al. 2005). Strong expression of both TC-Ascl1/E47 and TC-Brn2 can be seen in interphase nucleus following this labelling method (Figure 3B-C). In mitotic cells, TC-Ascl1/E47 is found excluded from mitotic chromatin, while TC-Brn2 colocalizes with DNA (Figure 3B-D). Thus, use of TC-tagged TFs validates previous observations, and indicates that the size of eGFP is not the cause of

**Figure 3.**
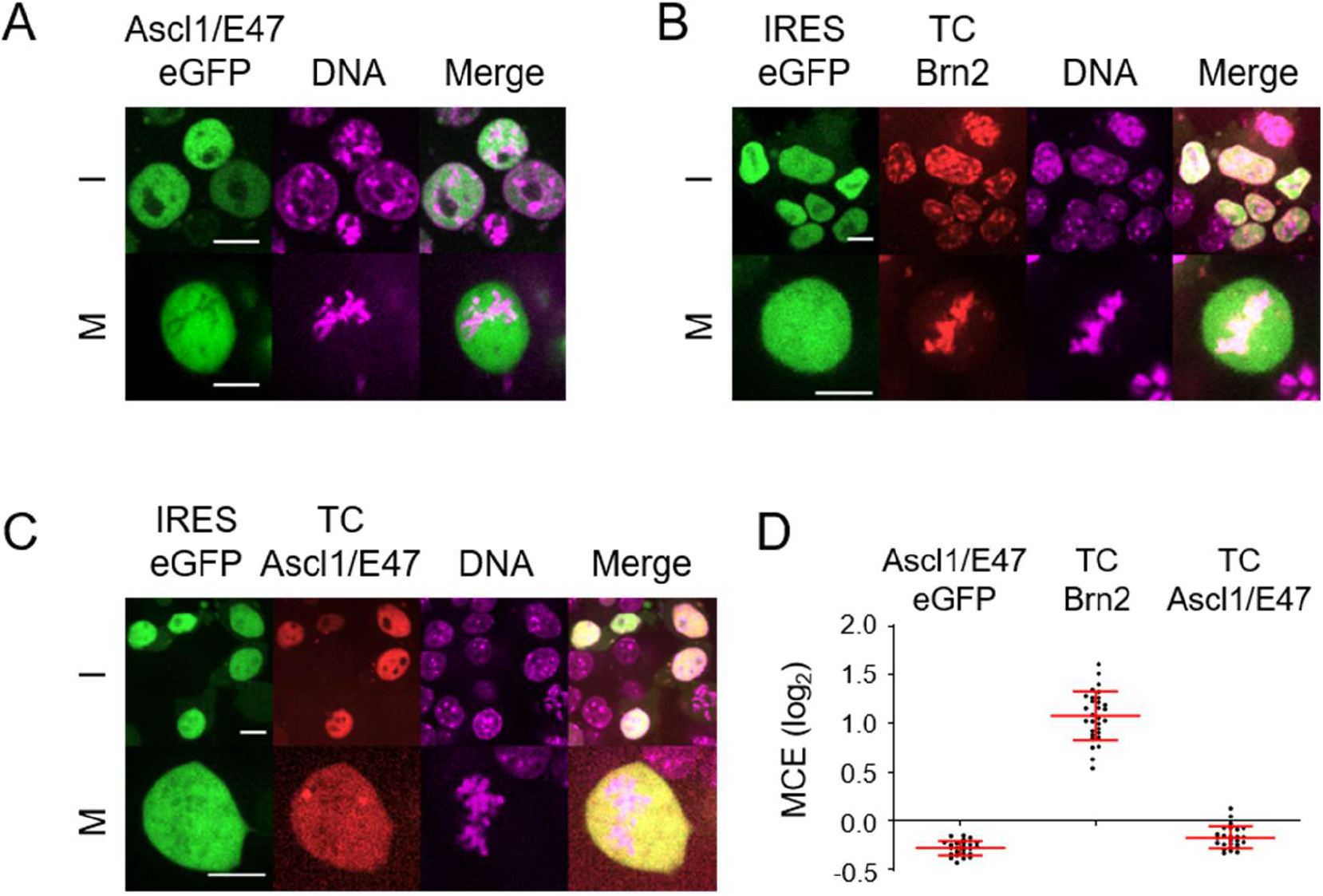
Tetracysteine-tag based live-cell imaging confirms distinct abilities of Ascl1 and Brn2 to interact with mitotic chromosomes. **(A-C)** Representative captures from live-cell imaging of P19 cells expressing Ascl1/E47-eGFP (A), TC-Brn2-IRES-GFP (B) or TC-Ascl1/E47-IRES-GFP (C) in presence of DNA dye SiR-Hoechst. **(D)** Quantifications of mitotic chromosome enrichment levels of all three conditions, measured using cells undergoing metaphase (n=25 in Ascl1/E47-eGFP; n=30 in TC-Brn2; n=26 in TC-Ascl1/E47). Data shown as mean ± SD. I=Interphase; M=Mitosis. Scale bars=10μm.

Ascl1 exclusion from mitotic chromatin. Moreover, Ascl1-eGFP was found to be transcriptionally active as judged by its ability to induce neuronal differentiation when expressed in P19 cells, or to activate reporter gene expression in transcriptional assays (Supplemental Figure S3). Altogether, these various lines of evidence indicate Ascl1 does not bind mitotic chromatin. Given previous studies have shown Ascl1 has pioneer TF activity (Wapinski et al. 2013; Raposo et al. 2015; Fernandez Garcia et al. 2019), our results clearly indicate these two activities (mitotic chromosome binding and pioneer) can be dissociated and may be deployed differently in reprogramming versus NS cell self-renewal.

### The DNA binding domain of Brn2 is both required and sufficient for mitotic chromosome binding

Next, we characterized the structural determinants of mitotic chromosome binding of Brn2, using live-cell imaging in transfected P19 cells. Using a truncated Brn2 derivative, we found its structurally conserved DNA-binding domain (DBD), composed of both the POU-specific (POUS) and the POU-homeo (POUH) domains, to be sufficient on its own to associate with mitotic chromosomes (Figure 4A-C). To further address the contribution of the Brn2 DBD, double point mutations were introduced in residues of either sub-domain, which mediate direct contact with DNA as assessed by structural data or *in vitro* binding assays (Malik, Zimmer, and Jauch 2018). These completely abolish (C311A, R312E) or severely reduce (N406Q, R407G) mitotic chromatin binding (mean MCE=-0.06 and 0.25, respectively), indicating the integrity of both POUS and POUH domains are required (Figure 4A-C). Of note, although the latter mutant retains some degree of mitotic chromosome binding, this becomes restricted to centromeric and pericentromeric regions (Figure 4B). Importantly, for each Brn2 derivative analyzed, the heterogeneous expression levels characteristic of transient transfection did not result in significant differences in MCE values across cells (Supplemental Figure S4A), highlighting the plausibility of our comparisons. In conclusion, the DBD is both required and sufficient to mediate the interaction of Brn2 with condensed chromosomes. Mitotic chromosome binding is likely to be a common feature of POU3F family members, according to live-cell imaging experiments in the context of full length Oct6 (aka POU3f1) or DBD-only (Oct6; Brn4 - aka POU3f4) derivatives (Supplemental Figure S4B-C).

**Figure 4.**
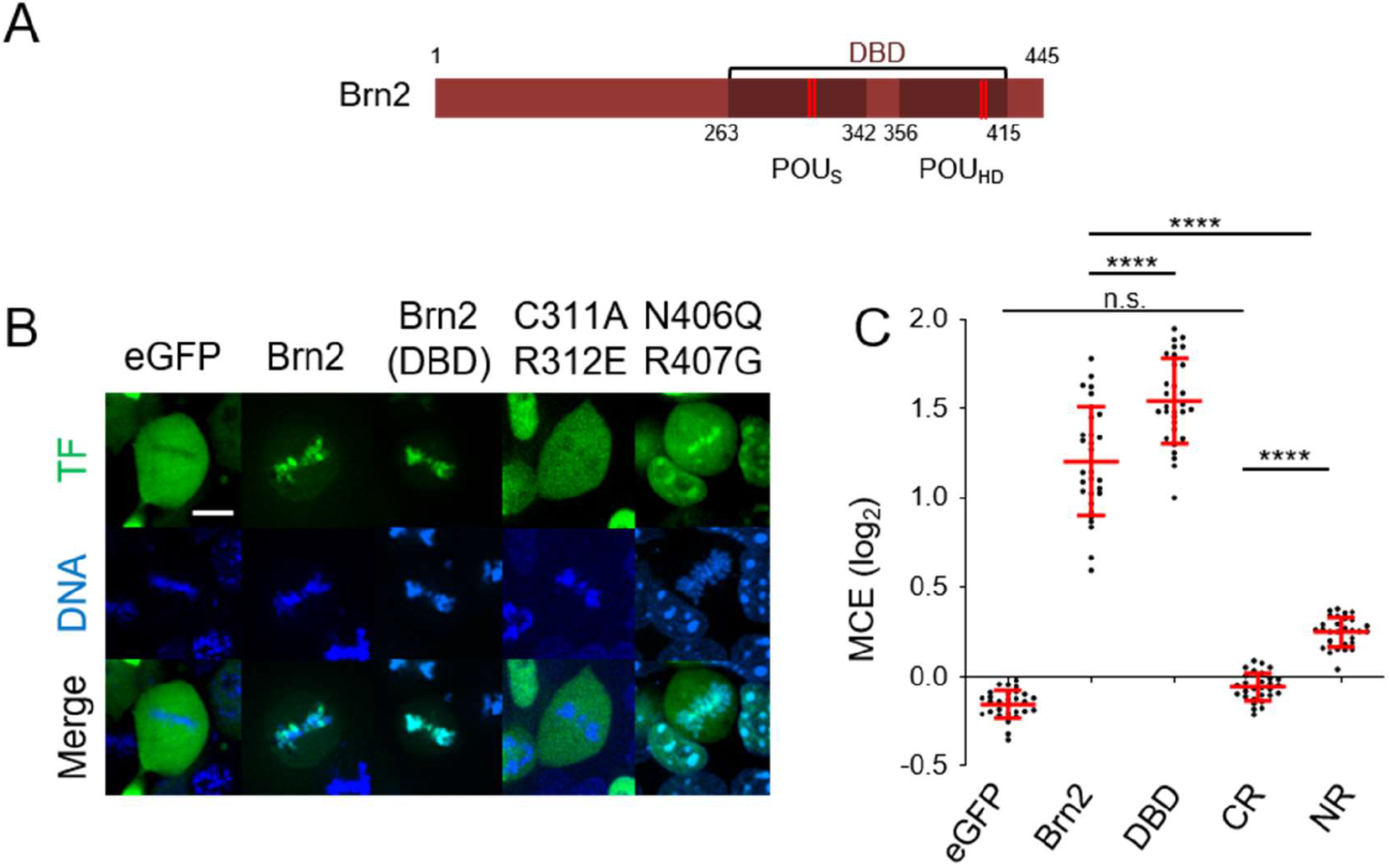
The DNA binding domain (DBD) of Brn2 is both required and sufficient for mitotic chromosome binding. **(A)** Representation of Brn2 protein domains to scale, including the DBD with its POUs and POUh domains. Red stripes mark residues involved in sequence-specific binding that were mutated in live imaging experiments. **(B)** Representative captures from live-cell imaging of P19 cells expressing eGFP fusion proteins of full-length Brn2, its truncated DBD or Brn2 derivatives with mutations in residues within the POUs domain (C311A/R312E) or POUh domain (N406Q/R407G). Control eGFP is also shown. Cells were synchronized in metaphase using proTAME and Apcin and live imaged together with DNA staining Hoechst. **(C)** Quantifications of mitotic chromosome enrichment levels from different Brn2 derivatives. Data shown as mean ± SD (n=30 for each condition). One-way ANOVA Tukey’s multiple comparison test was performed. p>0.05 (n.s.), p≤0.05 (*), p≤0.01 (**), p≤0.001 (***), p≤0.0001 (****). Scale bar = 10 μm.

### Brn2 does not establish sequence-specific interactions with mitotic chromatin

In order to map the genomic regions occupied by Brn2 in mitosis, we compared the genome-wide binding profile of endogenous Brn2 in NS cells in asynchronous (interphase) and mitotically arrested NS cells, using chromatin immunoprecipitation followed by deep sequencing (ChIP-seq). Colchicine incubation followed by shake-off resulted in a mitotic population of NS cells highly enriched in prometaphase (>90%), while maintaining mitotic chromosome binding in live-cell imaging, and similar Brn2 expression levels as assessed by western-blot analysis (Supplemental Figures S5A-B and S6A-B). Location analysis in interphase identified a high number (6,770) of Brn2 binding events, including within previously characterized neural enhancers driving expression of Nestin and Fabp7 genes (Figure 5A-C). Gene ontology analysis of genes associated with Brn2 binding showed enrichment for terms related with its proposed function in NS cells, including “somatic-” and “neuronal stem cell maintenance” (Supplemental Figure S5E). A *de novo* search for enriched DNA motifs present at peak summits found MORE and octamer motifs as main mediators of Brn2 binding (as dimer and monomer, respectively), with a bin-by-bin analysis indicating the MORE motif to be prevalent in the top 1,000 peaks (Figure 5D). Strikingly, sequence-specific binding is drastically reduced in mitosis, with only 85 binding events identified corresponding to genomic regions associated with strong Brn2 binding in interphase (Figure 5A-C). Moreover, no mitotic specific peaks were identified, altogether suggesting the <2% peaks that remain in the mitotic sample may result from contamination with interphase cells. Absence of sequence-specific binding is also in line with the observation that a mutation previously described to abolish Brn2 homodimerization (M414N), a required condition for binding to the MORE motif, does not decrease Brn2 mitotic chromosome binding (Figure 5E-F) (Dugast-Darzacq, Egloff, and Weber 2004; Jerabek et al. 2017).

**Figure 5.**
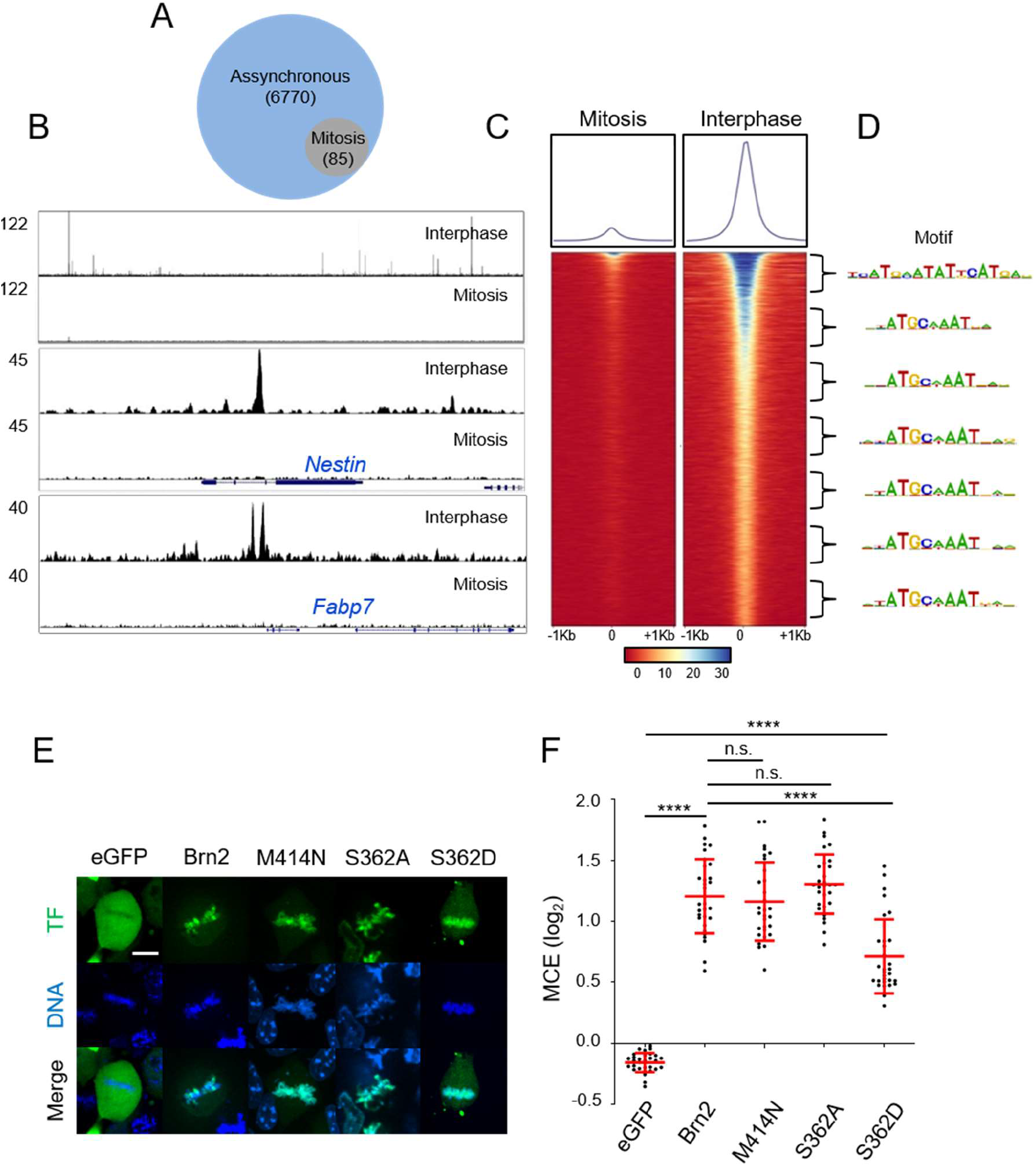
Brn2 mitotic chromosome binding is not mediated by sequence-specific interactions. **(A)** Venn diagram depicting the number of genomic regions bound by Brn2 in mitotically synchronized (95% purity sample) or unsynchronized cultures, as determined by ChIP-seq. **(B)** ChIP-seq Brn2 binding profile in a large region of chromosome twelve showing multiple peaks found in unsynchronized sample, but no binding in mitosis (top panel). Brn2 binding was found at expected regions within previously characterized neural enhancers in Nestin and Fabp7 genes, only in the interphase sample (middle and bottom panels). **(C)** Density plot of ChIP-seq reads from mitotic and interphase samples, mapping to genomic regions centered on Brn2 peak summits found in interphase. Signal intensity represents normalized tag count, ordered by increasing p-values. **(D)** A bin-by-bin search for enriched DNA binding motifs centered at Brn2 peak summits identifies the MORE motif as the highest enriched on the top 1000 peaks, while the octamer motif is most enriched in other bins. **(E)** Representative captures from live-cell imaging of P19 cells expressing eGFP fusion proteins of full length Brn2, a Brn2 mutant that cannot homodimerize (M414N), and a phospho-dead (S362A) or phospho-mimetic (S262D) Brn2 derivative in a residue targeted by mitotic phosphorylation that hampers sequence-specific binding. Control eGFP is also shown. Cells were synchronized in metaphase using proTAME and Apcin and live imaged together with DNA staining Hoechst. **(D)** Quantifications of mitotic chromosome enrichment levels from live imaging experiments shown in (E). Data shown as mean ± SD (n=30 for each condition). One-way ANOVA Tukey’s multiple comparison test was performed. p>0.05 (n.s.), p≤0.05 (*), p≤0.01 (**), p≤0.001 (***), p≤0.0001 (****). Scale bar = 10 μm.

### Phosphorylation of Ser362 impairs Brn2 sequence-specific interactions but is compatible with mitotic chromosome binding

Post-translational modifications of Brn2 during mitosis, in particular phosphorylation, could account for its lack of sequence-specific binding. Western blot analysis revealed a Brn2 specific band more diffused in the mitotic protein extracts (which could be reverted upon phosphatase treatment), confirming Brn2 is a phospho-protein in mitotic NS cells (Supplemental Figure S6A-B). Of notice, phosphorylation of a conserved serine (Ser362 in Brn2) that inhibits sequence specific DNA-binding of POU proteins was previously shown to occur during G2/M in multiple cell types (Nieto et al. 2007; Sunabori et al. 2008; Saxe et al. 2009; Jang et al. 2012). To test whether Brn2 phosphorylation could indeed abolish sequence-specific binding without compromising mitotic chromosome association, we tested the behavior of a phospho-mimetic derivative (Brn2 S362D). Fluorescence Recovery After Photobleaching (FRAP) assays reveal that Brn2 S362D displayed similar recovery curves to the Brn2 DBD mutant C311A/R312E, in line with *in vitro* studies showing Brn2 S362D is devoid of sequence-specific binding (Supplemental Figure S6C-E). Furthermore, this phospho-mimetic form of Brn2 retains a significant ability to associate with mitotic chromosomes (MCE= 0,7 vs 1,2 for WT Brn2 vs 1,3 for its phospho-dead mutant, S362A) (Figure 5E-F). Thus, mitotic specific phosphorylation of Brn2 (in particular serine 362) provides a possible molecular basis that conciliates mitotic chromosome binding with the absence of sequence-specific interactions.

### Differences in Brn2 and Ascl1 mitotic chromatin binding ability derive from their distinct electrostatic properties

Electrostatic interactions have been proposed to play a major role in mediating the association between TFs and highly compacted DNA (Raccaud et al. 2019). To better understand the contribution of electrostatic forces to the interaction of Brn2 and Ascl1 with mitotic chromatin, we extended our live-cell imaging analysis to cells in interphase. The colocalization of each TF with DNA was characterized by partitioning the cell nucleus in regions of various chromatin densities (heterochromatic, DNA-rich and DNA-poor), using Hoechst staining as previously described (Raccaud et al. 2019) (Figure 6-A). In NS cells, Brn2 was found highly enriched in dense and heterochromatic regions, a property known to correlate with mitotic chromatin binding (Figure 6B and D) (Raccaud et al. 2019). Depletion of Brn2 from DNA-poor regions was also observed in transfected P19 cells. Interestingly, S362D Brn2 (which is devoid of sequence-specific binding but binds mitotic chromatin) presented the same distribution as Brn2, whereas C311A/R312E Brn2 (unable to bind mitotic chromosomes) was evenly distributed across the three domains (Figure 6C-D).

**Figure 6.**
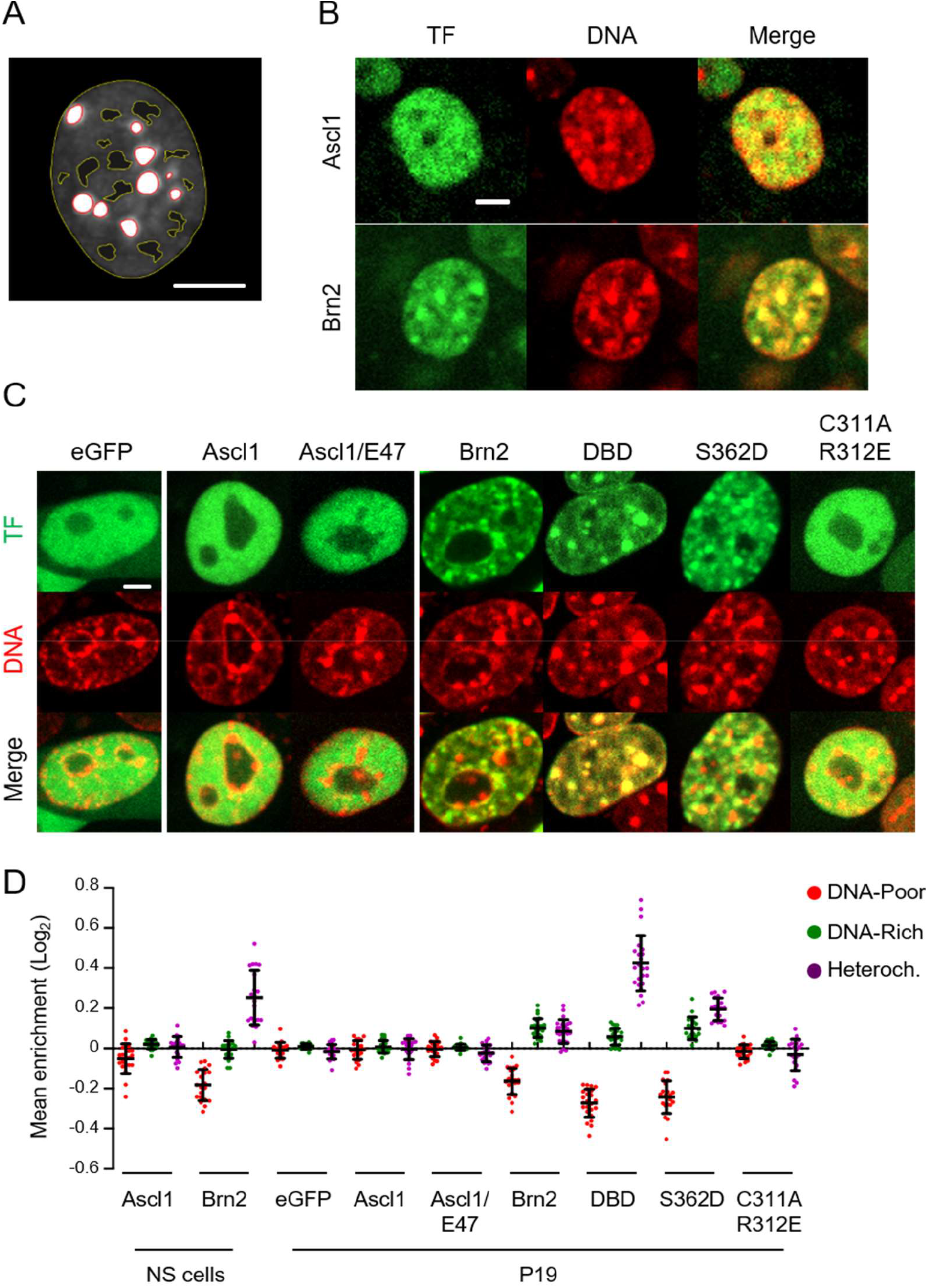
Brn2 (but not Ascl1) associates with highly compacted DNA regions in the interphase nucleus. **(A)** Image depicting the segmentation of an NS cell nucleus into heterochromatic DNA regions (delineated in red), DNA-poor regions (delineated in yellow) and DNA-rich regions (space in between). **(B)** Representative captures from live cell-imaging of eGFP-tagged Ascl1 and Brn2 expressing NS cells, stained with Hoechst for segmentation into nuclear regions with different chromatin densities. **(C)** Representative captures from live-cell imaging of P19 cells expressing variants of Ascl1 and Brn2 eGFP fusion proteins as indicated in figure, stained with Hoechst for segmentation into nuclear regions with different chromatin densities. **(D)** Quantification of imaging analyzes shown in (B) and (C). Enrichment levels at different segmented DNA regions are shown as mean ± SD. (n=20 cells for most conditions; n=24 cells for Brn2 and DBD both in P19 cells). Scale bars=5μm.

In sharp contrast to Brn2, Ascl1 was not found preferentially at heterochromatic or DNA-dense regions, when compared to DNA-poor regions, in either cell type (Figure 6B-D). This did not result from an excess of Ascl1 protein over its heterodimeric partners, as a similar result was obtained with Ascl1/E47 tether (Figure 6C-D). Thus, our results indicate that Brn2 and Ascl1 have very different abilities to associate with highly compacted DNA, indicating the differences observed in mitotic chromosome binding rely on their different electrostatic properties. To further confirm this, we sought to change the electrostatic potential of Ascl1 and assess its impact on the association with mitotic chromatin. Fusion of three highly positively charged SV40 NLS sequences promotes eGFP association with mitotic chromosomes, as assessed by live-cell imaging (mean MCE=0.37) (Supplemental Figure S7). This is known to result from an increase in electrostatic interactions, independently of the canonical NLS function (Raccaud et al. 2019). Strikingly, the observed increase in MCE was strongly counteracted by the addition of Ascl1 in fusion with eGFP, even when in the context of an obligatory dimer with E47 (mean MCE=0.01 and 0.08, respectively). A nearly full recovery of MCE levels was only attained upon fusion of additional NLS sequences (mean MCE=0.31) (Supplemental Figure S7), altogether in line with a low electrostatic potential of Ascl1. In conclusion, our results from live-cell imaging in both interphase and mitosis underlie differences of electrostatic properties of Brn2 and Ascl1, and are in line with non-specific electrostatic interactions being the main determinants for the association of these TFs with mitotic DNA.

### Targets of Brn2 (Nestin) and Ascl1 (Dll1) have distinct onsets of reactivation during M-G1

A mitotic bookmarking function entails that gene regulatory information is conveyed by TF binding to specific gene regulatory regions. Even in the absence of sequence-specific interactions, retention of Brn2 on mitotic chromatin may increase its local concentration, thereby facilitating how it searches and reactivates target genes during the transition from mitosis-to-G1 (M-G1). Recent studies suggest the reactivation of a fraction of the transcriptome starts early during M-G1 transition (i.e. late metaphase) (Palozola et al. 2017; Zhang et al. 2019). Thus, Ascl1 dependency on nuclear import may result in a delayed onset of reactivation of its transcriptional program, as compared to Brn2. Proceeding with testing our hypothesis, we next investigated the kinetics of transcriptional reactivation of genes encoding the intermediate filament Nestin and the Notch ligand Dll1, two well-established targets directly activated by Brn2 and Ascl1, respectively, suggested by multiple studies to function as readout of these TFs. The requirement of Brn2 for efficient activity of the Nestin neural enhancer was thoroughly documented (Josephson et al. 1998; Sunabori et al. 2008; Lodato et al. 2013). The strict dependency of Dll1 on Ascl1, together with its pattern and kinetics of expression, makes Dll1 a good proxy for Ascl1 activity in neural progenitor cells (Castro et al. 2006; Kageyama et al. 2008; Shimojo et al. 2016).

In order to evaluate the dynamics of transcription during M-G1 transition, we used single molecule fluorescence in situ hybridization (smRNA-FISH) to simultaneously quantify nascent and mature mRNA in single asynchronous NS cells, using spectrally distinguishable probes against introns or exons. In all interphase NS cells analyzed, exonic signal was much more prevalent than the intronic signal, as expected given the high relative abundance of mature transcripts (Figure 7A and C). Although many exon foci were found in each interphase cell, particularly for Nestin probe, colocalization of exonic and intronic probes on DNA (marking primary transcripts) was found to be proportional to the number of loci (0-4 depending if in G1 or G2 phase) (Figure 7A and C). Colocalizing foci from primary transcripts, a readout of active transcription, were often the brightest to be found in the cell. As expected, exon and intron probe signals almost completely disappeared in interphase cells upon RNase treatment (Figure 7E-F).

**Figure 7.**
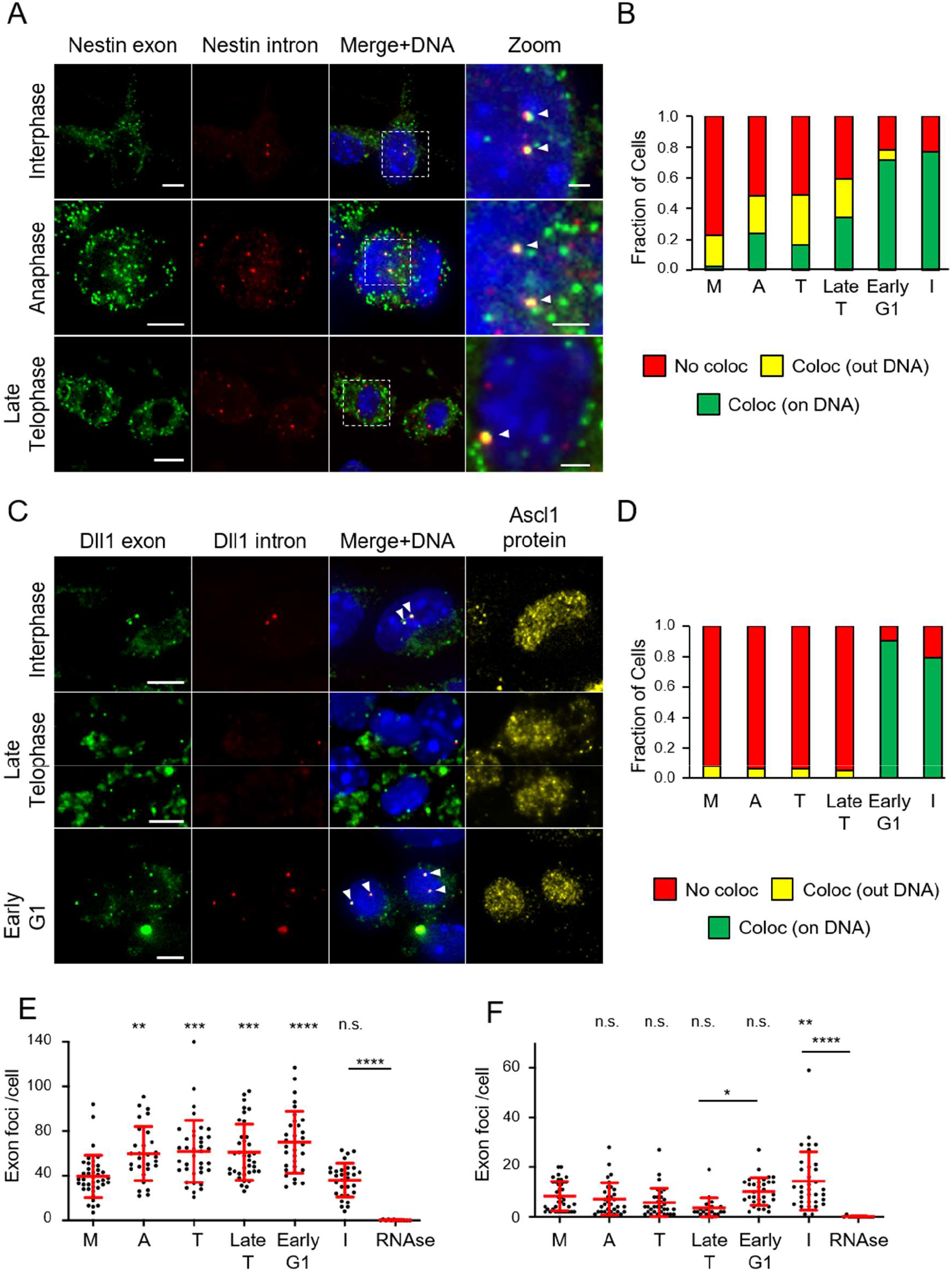
Targets of Brn2 (Nestin) and Ascl1 (Dll1) show distinct kinetics of transcriptional reactivation during M-G1 transition in proliferating NS cells. **(A)** Representative images of cells in interphase, anaphase and late telophase stained by smRNA-FISH using exonic (FAM) and intronic (Q570) probes for Nestin transcript. Images shown are maximum intensity projections of six optical planes with 0,3μm Z step intervals, with white arrow-heads marking spots of colocalized intron-exon probe signal found on DNA (stained with DAPI) Scale bars=5μm. **(B)** Stacked bar plots showing the fractions of cells containing at least one spot of colocalized intron-exon probe signal found on DNA (green), outside DNA (yellow) or without colocalization spots (red) (n=40 in metaphase; n=29 in anaphase; n=37 in telophase, n=37 in late telophase; n=32 in early G1; n=30 in interphase; n=12 in interphase + RNase). **(C)** Representative images of cells in interphase, late telophase and early G1 co-stained by smRNA-FISH using exonic (FAM) and intronic (Q570) probes for Dll1 transcript, and by immuno-cytochemistry for Ascl1 protein. Images shown are maximum intensity projections of six optical planes with 0,3μm Z step intervals, with white arrow-heads marking spots of colocalized intron-exon probe signal found on DNA (stained with DAPI). Scale bars=5μm. **(D)** Stacked bar plots showing the fraction of cells containing at least one spot of colocalized intron/exon probe signal found on DNA (green), outside DNA (yellow) or without colocalization spots (red) (n=32 for metaphase, anaphase, telophase, early G1; n=21 for late telophase; n=34 for interphase; n=11 for interphase + RNase). **(E-F)** Quantifications of exon probe signal per cell for Nestin (E) and Dll1 (F) transcripts at different stages during MG1 transition, in experiments described in (A-D). Quantification of RNAse treated sample is included as control. One-way ANOVA Tukey’s multiple comparison test was performed. p>0.05 (n.s.), p≤0.05 (*), p≤0.01 (**), p≤0.001 (***), p≤0.0001 (****). M=metaphase; A=anaphase; T=telophase; Late T=late telophase; I=interphase. Data shown as mean ± SD.

When assessing transcription during M-G1 transition, we restricted the analysis to Ascl1 expressing cells, with its cellular distribution (assessed by co-immunostaining) used as a proxy for the timing of nuclear envelope reformation. Telophase cells were defined by having cytokinesis undergoing prior to Ascl1 import, whereas cells in the immediate subsequent step (when Ascl1 import could already be observed) were considered in late telophase (Figure 7C). In the case of Nestin, a significant fraction of actively transcribing cells was found to increase steadily throughout M-G1 transition, starting as early as anaphase (7/29 cells; 0,24) and until early G1, when the largest number of cells with intron/exon colocalization on DNA was observed (23/32 cells; 0,72) (Figure 7A-B). Active transcription was often observed only in one daughter cell, or in both but in distinct locations, recalling recently found heterogeneity in allele location and 3D interactions at single-cell level during interphase (Finn et al. 2019). At all stages during M-G1 transition, a significant number of intron/exon colocalized foci was observed outside DNA, reaching the highest fraction in telophase (12/37 cells; 0,32) (Figure 7A-B). However, these were almost never found in interphase, and are likely to represent long-lived unprocessed transcripts resulting from some level of RNA splicing inhibition during mitosis (C. Shin and Manley 2002), and which diffused away from the transcription site. Moreover, the presence of active Nestin transcription during M-G1 transition is also supported by a concomitant increase in total number of transcripts (exon probe foci) per cell (Figure 7E). As expected, intron/exon probe colocalization for Nestin transcript almost disappeared upon incubation with transcriptional inhibitor triptolide (Supplemental Figure S8A-B). This was concomitant with a strong reduction of intron signal (from an average of 10 to 3 foci per cell), while no increase in exon probe counting was observed throughout M-G1, altogether in line with transcription occurring during this period in control conditions (Supplemental Figure S8C-D).

By contrast to the Nestin gene, analysis of Ascl1-dependent Dll1 transcription revealed absence of actively transcribing cells until late Telophase, with a dramatic increase in colocalization of intronic and exonic probes observed in early G1 (29/32 cells; 0,91) (Figure 7C-D). Quantification of exonic probe foci number revealed no increase during M-G1 transition, in line with Dll1 transcription only restarting in early G1 (Figure 7F). Finally, we have repeated the same analysis using probes for Nestin and Dll1 transcripts labelled with a different set of fluorophores, and obtained similar results when quantifying exon/intron colocalization events (Supplemental Figure S9). In conclusion, surveying active transcription with smRNA-FISH shows the onset of activation of Nestin and Dll1 genes occurs at distinct timepoints during M-G1 transition.

### Reactivation of Nestin transcription in early M-G1 is dependent on Brn2

We next sought to understand whether transcriptional reactivation of Nestin during early stages of M-G1 transition depends on Brn2 function. To address this, a mitotic-specific dominant negative version of this TF (DN-Brn2) was generated, by fusing nuclear export signal (NES) sequences at each flank of the Brn2 DBD, fused to a fluorescent reporter protein (mCherry). This strategy was used to force the localization of this dominant form outside the nucleus throughout interphase, gaining access to chromatin only during mitosis, upon NEBD.

Expression of DN-Brn2 was placed under the control of a tetracycline inducible promoter, in NS cells where endogenous Brn2 was tagged with eGFP (Figure 8A-B). Upon treatment with the tetracycline analogue doxycycline (Dox), expression of DN-Brn2 is found in association with mitotic chromosomes, with its rapid export observed upon nuclear envelope reformation at the end of M-G1 transition (80% exported in 20 minutes after telophase) (Figure 8C). The association of DN-Brn2 with condensed chromosomes during metaphase did not alter the association of Brn2-eGFP (MCE=0.42 vs 0.46 with or without Dox respectively, with R^2^=0.1175) (Figure 8D-E). This result implies that DN-Brn2 is unable to compete and prevent Brn2-eGFP association with metaphase chromosomes. This finding confirms that binding to mitotic chromatin (before the onset of mitotic exit) does not rely on the limited number of sequence-specific Brn2 sites available in the genome. Accordingly, we detected a linear increase in MCE levels of Brn2, observed in overexpression experiments (Supplemental Figure S4A). Yet, such DN-Brn2 is expected to compete with endogenous Brn2 once sequence-specific transcription is initiated (early anaphase, see Figure 7A-B). To understand whether reactivation of Nestin transcription during anaphase depends on Brn2 function, we compared the levels of transcripts (using smRNA-FISH) in the presence or absence of DN-Brn2. Strikingly, a significant reduction of Nestin transcripts was observed in the presence of DN-Brn2, as indicated by a decrease in the number of exon probe foci in Dox treated cells, and by their levels remaining unchanged until late telophase (Figure 8F-G). This finding confirms the requirement of the canonical TF activity of Brn2 in reactivation of Nestin gene during early stages of M-G1 transition.

**Figure 8.**
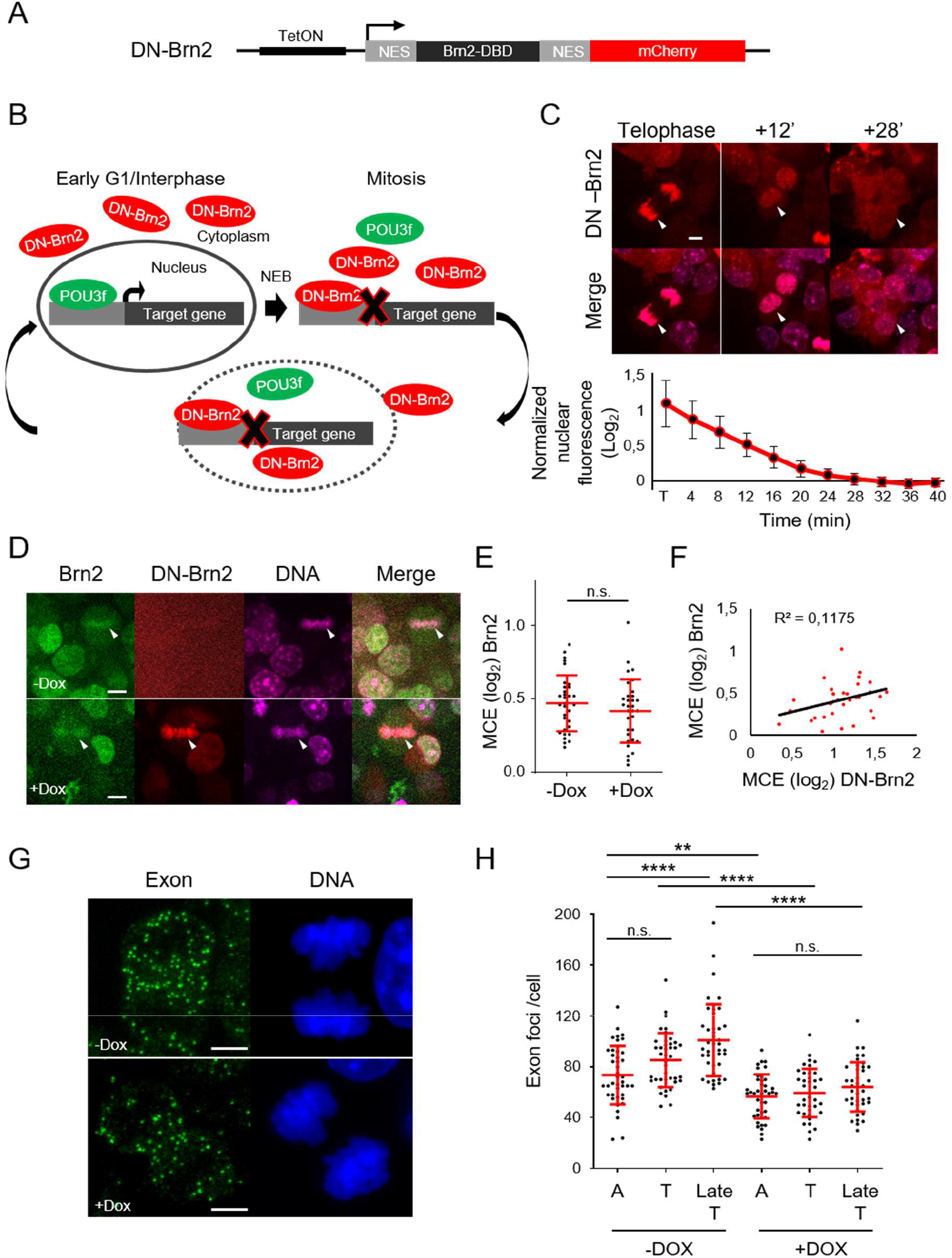
Transcription reactivation of Nestin gene during M-G1 transition is dependent on Brn2 activity. **(A)** Experimental strategy whereby a mitotic-specific dominant-negative form of Brn2 (DN-Brn2) composed of its DBD flanked by nuclear export signal (NES) sequences and in fusion with mCherry, was expressed in NS cells under a Dox-inducible promoter. **(B)** Schematic depicting how DN-Brn2 competes with POU3 transcription factor family members in cell-cycle stages with or devoid of nuclear envelope. **(C)** Time-lapse live-cell imaging of Dox treated cells expressing DN-Brn2, from telophase until G1. White arrow-heads show association of DN-Brn2 with mitotic DNA (labeled with SiR-hoechst) during telophase, and its cytoplasmic localization after nuclear envelope reformation. Quantification of nuclear export kinetics of dN-Brn2, normalized to interphase levels, is shown below. n=11 cells. Scale bar=5μm **(D)** Representative captions from live-cell imaging of DN-Brn2-NS cells in the absence (top) or presence (bottom) of Dox. White arrow-heads indicate an example of a metaphase plate in each condition. DNA labeled with SiR-hoechst. Scale bars=5μm **(E)** Quantification of mitotic chromosome enrichment levels of Brn2-eGFP with or without Dox treatment reveals no significant difference between conditions. Mann-Whitney test was performed; p>0.05 (n.s.). n=32 for each condition. **(F)** Representative images of Dox treated and untreated NS cells in telophase stained by smRNA-FISH using exonic (Q670) probe for Nestin transcript. DNA staining with DAPI. Images shown are maximum intensity projections of twelve optical planes with 0,3μm Z step intervals. Scale bars=5μm **(G)** Quantifications of exon probe signal per cell for Nestin transcript at different stages during MG1 transition. One-way ANOVA Tukey’s multiple comparison test was performed. p>0.05 (n.s.), p≤0.05 (*), p≤0.01 (**), p≤0.001 (***), p≤0.0001 (****). n=41 for all untreated cells (-Dox). n=41 for A with Dox, 38 Tel with Dox and 41 Late Tel with Dox. A=anaphase; T=telophase; Late T=late telophase; Data shown as mean ± SD.

## Discussion

Contrary to the classical view of mitotic chromatin as being transcriptional inert and refractory to TF binding, various studies have in recent years described the ability of certain TFs to associate with mitotic chromatin in dividing cells. However, the molecular basis and functional consequences of such interactions have remained the subject of controversy, with the concept of “mitotic bookmarking by TFs” at the center of it. Recently, genome wide profiling of transcription through each stage of mitosis revealed that reactivation of the transcriptome occurs in temporally coordinated waves starting early during M-G1 transition (Palozola et al. 2017). These observations raise the question of what is the underlying regulatory logic during this period, and how mitotic bookmarking by TFs may be involved in this process. In the present work, we have tackled this question in neural stem cells, a cell type which can be experimentally manipulated and where key TFs are known. Focusing on Brn2 and Ascl1, we investigated how two TFs with distinct functions in neural cell identity interact with mitotic chromatin, and how this impacts their transcriptional output during M-G1 transition.

Important mechanistic insights into the temporal pattern of gene reactivation during M-G1 transition have recently started to emerge. Enhancer usage, and the establishment of enhancer-promoter loops, has been observed as early as anaphase (Palozola et al. 2017; Zhang et al. 2019; Abramo et al. 2019). A role for TFs in regulating chromatin accessibility during this period has been recently shown (Friman et al. 2019; Owens et al. 2019; Festuccia et al. 2019); however, a clear link between canonical TF activity and timing of gene reactivation has remained elusive. Addressing this requires the ability to knock-down the function of TFs specifically during mitosis, a task made more difficult with Brn2, by the co-expression of redundant POU3f family members in neural stem/progenitor cells (Sugitani et al. 2002; McEvilly et al. 2002). Using a mitotic-specific dominant-negative approach we were able to overcome these limitations, providing evidence for the need of POU3f TFs in Nestin gene transcription starting from anaphase. This was made possible also by the use of single-cell smRNA-FISH, which provided the sensitivity and temporal resolution that sets aside our study from previous reports using cell-population analysis. Although we cannot exclude that other Brn2 targets may have later onsets of expression due to other regulatory constraints, here we showed that the transcriptional activity of Brn2 is non-limiting from the start of mitotic exit.

The concept of mitotic “bookmarking” by TFs as a mechanism to convey gene regulatory information across cell division, implies the establishment of sequence-specific interactions with regulatory regions throughout the duration of mitosis/mitotic exit. Our ChIP-seq analysis revealed absence of sequence-specific binding by Brn2 in cells arrested in prometaphase. We propose phosphorylation of Ser362 as a possible mechanism to impair sequence-specific binding. The equivalent residue in Oct4 is a target of Aurora B kinase during prometaphase, being subsequently dephosphorylated by active protein phosphatase 1 (PP1) during mitotic exit (J. Shin et al. 2016; Cai et al. 2018). Notably, the docking site for PP1 is also conserved in Brn2, with the timing of PP1 activity being compatible with dephosphorylation of Brn2 on time for activation of Nestin gene transcription early in M-G1. In line with these results, we virtually did not detect any ongoing Nestin transcription during metaphase. This suggests reactivation of Nestin does not require the maintenance of low levels transcription throughout mitosis that was recently observed for some genes (Palozola et al. 2017).

The absence of sequence-specific interactions observed with ChIP-seq is in apparent contradiction with mutagenesis of residues in helices 3 of each POUS and POUH sub-domains, predicted to establish base-specific interactions with an octamer motif, and which impair binding to mitotic chromosomes (Malik, Zimmer, and Jauch 2018). However, these observations can be conciliated, as TFs often rely on the DBD when undergoing electrostatic-guided one-dimensional (1D) search, and the same residues can switch role from a purely electrostatic interaction with the DNA backbone, to a highly specific binding mode (Kalodimos et al. 2004; Suter 2020). Thus, mutagenesis experiments may not always be able to disentangle the different nature of TF-DNA interactions.

We found Brn2 association with mitotic chromatin to be governed mostly by non-specific electrostatic interactions. This important observation indicates Brn2 does not function as a bona-fide mitotic bookmarker in neural stem/progenitor cells. As an alternative, we propose that association of Brn2 with mitotic chromosomes may serve to increase its local concentration near chromatin during M-G1 transition, thereby facilitating its search for target genes. This model was indeed supported by the early timing of reactivation of Nestin transcription during mitotic exit, in a Brn2 dependent manner. By contrast, exclusion of Ascl1 from mitotic chromatin is associated with a late reactivation of its target Dll1 in early G1. This occurs concomitant with a sharp increase of Ascl1 concentration near chromatin, which we show requires active nuclear import of TF into the newly reformed nuclear envelope (see model in Figure 9). Our results support the emerging view that TFs association with mitotic chromosomes detected by live-cell imaging is to large extent driven by non-specific interactions (Festuccia et al. 2019; Raccaud et al. 2019). However, to which extent the lack of sequence-specific binding observed with Brn2 can be extended to other mitotic chromosome binding TFs, must be examined in future. Of note, the Brn2 case finds precedent in a recent study showing association of Sox2 with mitotic chromatin in ES cells also occurs in absence of sequence-specific binding (Festuccia et al. 2019).

**Figure 9.**
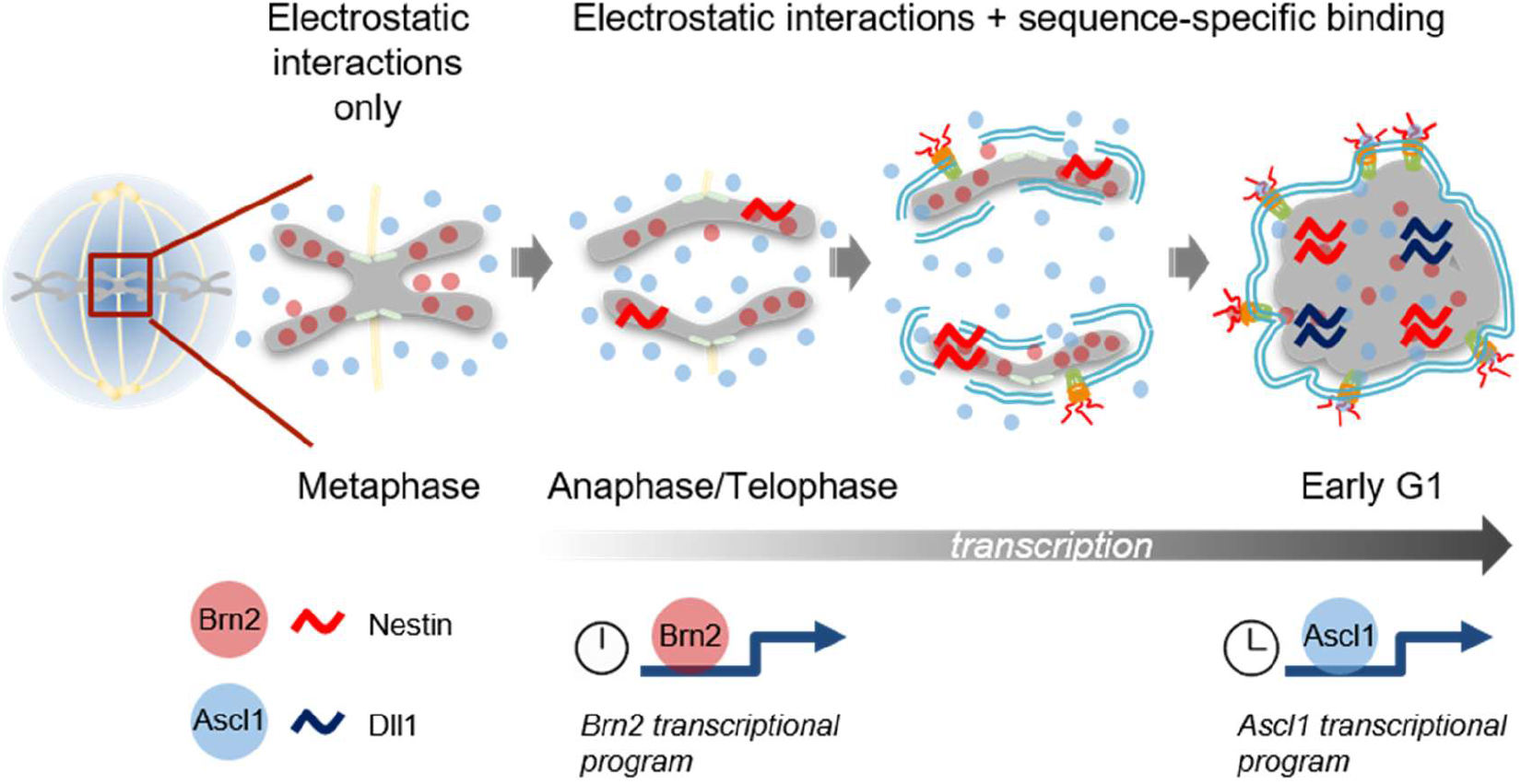
Model for hierarchical reactivation of transcription by Brn2 and Ascl1, in neural stem/progenitor cells undergoing proliferative divisions. Binding of Brn2 to metaphase chromosomes does not involve sequence-specific binding and is mediated by electrostatic interactions. Constant presence near chromatin results in Brn2 inducing the transcriptional reactivation of its target gene Nestin starting from anaphase, a process which requires sequence-specific binding and occurs concomitantly with chromatin decondensation and Brn2 dephosphorylation (events not represented in figure). By contrast, Ascl1 is excluded from mitotic chromatin, given its low electrostatic potential. Ascl1 concentration in the vicinity of chromatin increases due to nuclear import (upon nuclear envelope reformation), resulting in transcriptional reactivation of its target gene Dll1 in early G1. The different dynamics observed favors an early versus late reactivation of transcriptional targets of Brn2 and Ascl1, respectively.

In contrast to Brn2, Ascl1 exclusion from mitotic chromatin or DNA rich regions in interphase is suggestive of low electrostatic potential, further supported by its strong ability to counteract the colocalization of SV40 NLS sequences with condensed chromosomes. As a possible consequence, Brn2 and Ascl1 may use different mechanisms when searching the interphase genome for target genes. The presence of Brn2 in the vicinity of mitotic chromatin and interphase DNA-rich regions, is in line with Brn2 using more efficiently 1D search-based mechanisms to find its target genes. On the contrary, and given its low electrostatic potential, Ascl1 may use preferentially random encountering (3D search), a mechanism highly dependent on protein levels (Steffen and Ringrose 2014). This is supported by observations showing Ascl1 cannot induce Dll1 transcription in late telophase, before its highest concentration inside the newly reformed nuclear envelope is reached.

Studies using stem/progenitor cells (including of neural origin) have shown transition through mitosis and G1 phase is crucial for establishing a window of opportunity for executing cell fate decisions (McConnell and Kaznowski 1991; Soufi and Dalton 2016). Understanding the temporal dynamics of transcription during M-G1 transition should cast light into how TFs with a role in cell identity operate. Antagonistic cross-regulatory interactions between pathways promoting progenitor maintenance and neuronal differentiation are well documented. Thus, the intricate balance that regulates cell identity in daughter cells should rely on the kinetics that define how such programs are rewired during mitotic exit. The model we propose describes how the timing of transcriptional reactivation of Brn2 targets may be determined, favoring the res-establishment of the neural stem/progenitor program (Figure 9). In the case of Ascl1, its expression in neural stem/progenitor cells undergoing proliferative divisions is in apparent contradiction with a well-established ability to promote neuronal commitment and differentiation. The current view is that Ascl1 neurogenic activity is kept at reduced level in a proliferative cell context, by mechanisms that may include its (oscillatory) mode of expression and post-translational modifications. We suggest the observed exclusion of Ascl1 from mitotic chromosomes (and delayed transcriptional activity), as yet another layer of regulation to avert premature neuronal differentiation in cycling cells.

Overall, our work results in a model whereby TFs, and their electrostatic properties, play an important role in determining the timing of reactivation of target genes during M-G1 transition. Is such model, TFs occupy distinct hierarchical levels depending on how they interact with mitotic chromatin. It is tempting to speculate that TFs with bona-fide mitotic bookmarking function (i.e. maintain sequence-specific binding throughout mitosis) may be higher in this hierarchy. Emerging genome-wide studies mapping histone marks during M-G1 transition, should help dissecting further the role of TFs during this period (Hsiung et al. 2016; Javasky et al. 2018; Kang et al. 2020).

## Materials and Methods

### DNA constructs

Expression vectors for eGFP fusions were generated in eGFP N1 (Clontech, #6085-1) by subcloning of PCR fragments obtained using primers and template vectors shown in Supplemental Table S1. Mutations of specific residues were introduced by site-directed mutagenesis using oligonucleotides shown in Supplemental Table S2. Plasmids encoding NLS-fusion constructs were generated by annealing and sub-cloning oligonucleotides with complementary sequences (Supplemental Table S3) and BsrGI and XbaI cohesive ends into the C-terminal domain of either eGFP N1, Ascl1-eGFP or Ascl1/E47-eGFP. All other constructs are described in specific sections below. Constructs generated in this study were confirmed by Sanger sequencing. Full length Oct6-GFP, DBD (Oct6)-eGFP and DBD (Brn4)-eGFP expression plasmids were kindly provided by Michael Wegner (Institut fur Biochemie, University Erlangen-Nurnberg).

### Establishment, genome editing and culture of NS lines

NS cell cultures were established from E13.5 mouse ventral telencephalon as previously described (Conti et al. 2005) and maintained in mouse Neurocult basal media supplemented with proliferation supplement (Stem Cell Technologies, #05702,), 1% Penicillin-Streptomycin (ThermoFisher, #15140122), 10ng/ml EGF (Peprotech, #315-09), 10ng/ul bFGF (Peprotech, #100-18B) and 2μg/mL laminin (Sigma-Aldrich, #L2020). Cells were kept in T-flasks (Corning, #734-2705) and split every 2-3 days using accutase (Sigma-Aldrich, #A6964). For eGFP tagging of Brn2 and Ascl1 by CRISPR/Cas9, template vectors were generated by PCR amplification of both left and right homology arms of genomic regions and eGFP coding sequence, using primers from Supplemental Table S4, and cloned into pBluescript II KS (+/-) using Gibson Assembly (New England Biolabs, #174E2611S). For generation of vectors for expression of gRNAs, oligonucleotide pairs (CACCGCGTCCAGTGAACTCAAGCGG, AAACCCGCTTGAGTTCACTGGACGC for Brn2 and CACCGACTTTACCAACTGGTTCTG, AAACCAGAACCAGTTGGTAAAGTC for Ascl1) were cloned into pX330 (Cong et al. 2013) using BbsI. NS cells were electroporated using Neon™ nucleofection system. Briefly, cells were rinsed with PBS without Ca2+ and Mg2+ (Biowest, #L0615-500), dissociated with Accutase and neutralized with growth medium. Cells were centrifuged at 1000 × g for 5 minutes at RT, washed with PBS, centrifuged at 1000 × g for 5 minutes at RT and resuspended in Resuspension Buffer at a final density of 1,2 × 10^6^ cells/ 120 μl. For CRISPR/Cas9 experiments, 1ug of each plasmid was used in a ratio of 1:1. The cell-DNA mixture was gently mixed and aspirated into the Neon™ Tip (Life technologies, #MPK10025) which was inserted into the Neon™ Pipette and subject to an electric pulse (Pulse voltage: 1700; Pulse Width 20; Number of pulses: 1). Finally, cells were transferred to a 6 well plate containing pre-warmed complete Neurocult medium. Four days later, cultures were enriched for recombined cells (GFP+) using FACSorting and clones originated from single-cells allowed to expand in 96 well-plates. Success of genome editing was verified by PCR from genomic DNA and confirmed by western-blot analysis. Briefly, crude cell lysates were diluted in 2x Laemmli buffer (Sigma-Aldrich, #S3401) and denatured for 5 min at 95°C. Samples were separated in 12% SDS-PAGE gels and transferred to nitrocellulose membranes (GE Healthcare, #GE10600008) using standard procedures. Blots were probed with primary anti-Ascl1 (Abcam, #ab211327, 1:1000), anti-Brn2 (GeneTex, #GTX114650, 1:1000), anti-GAPDH (Cell Signaling, #2118, 1:1000) or anti-Histone3 (Cell Signaling, #4499, 1:1000) and HRP-conjugated rabbit secondary antibody (Jackson Immunoresearch, #111-035-144)

### Generation of DN-Brn2 expressing NS cells

DN-Brn2 expression vector was generated by PCR amplification of Brn2 DBD and addition of NES sequences using primers ATC ATC GAA TTC GAG AGT CAT GCT TCA ACT TCC TCC TCT TGA ACG CCT TAC CCT TGG AGG AGG AGG ACC GGG CCA CCC AGG CGC GCA C and GAT GAT GGA TCC CCA AGG GTA AGG CGT TCA AGA GGA GGA AGT TGA AGT CCT CCT CCT CCA CCC CCA TAC ACA TCC TCG GC and mBrn2 template plasmid (Sugitani et al. 2002) into mCherry N1 (Clontech). Subsequently, a transposase-mediated integration system was used. Subcloning of DN-Brn2 into pTRE3gIRESBSDDest (DN-pDest) was done using Gateway protocol (Thermo Fisher) using primers GGG GAC AAG TTT GTA CAA AAA AGC AGG CTT CAC CAT GCC GGG CCA CCC AGG CGC and GGG GAC CAC TTT GTA CAA GAA AGC TGG GTC CTA CTT GTA CAG CTC GTC CAT GCC GCC GGT GGA GTG G and DN-Brn2 expression vector as template. NS cells were nucleofected as above with plasmids DN-pDest, CAGTet3G and pBase at a ratio of 2:1:1. Clones were grown in 96-well plates upon dilution of nucleofected cells to <1 cell per well (confirmed by microscopy). Positive clones were selected using 5μg/mL blasticidin upon addition of 2μg/mL of doxycycline hyclate (Sigma-Aldrich, #D9891) to induce gene expression and antibiotic resistance. Duplicates of positive clones were tested for mCherry expression and further expanded.

### Live imaging and quantification of mitotic binding

P19 cell line was cultured in DMEM high glucose media (BioWest, #L0101) supplemented with 1% L-glutamine (ThermoFisher, #25030024), 1% Penicillin-Streptomycin (ThermoFisher, #15140122) and 10% Fetal Bovine Serum heat-inactivated (BioWest, #S181BH-500). For imaging analysis, P19 cells were transfected on 4 or 8-well polymer chambers (ibidi, #80426 and #80826, respectively) at 70% confluency, using Lipofectamine 2000 (Life Technologies, #11668-019) in the proportion of 1ug DNA to 3uL lipofectamine (1ug/well) according to manufacturer’s protocol. Prior to imaging, P19 cells were moved to phenol red-free DMEM/F12 medium supplemented with FBS (10x). Live-cell imaging of mitotic chromosomes of both NS and P19 cells was achieved by timelapse, or snapshot acquisition after cell synchronization in metaphase using 60μM proTAME (R&D systems, #I-440-01M) and 200μM Apcin (R&D systems, #I-444-05M) for 2-6 hours. Confocal Z-series stacks were acquired at 37°C and 5% CO^2^, on a Yokogawa CSU-X Spinning Disk confocal, using a 60x 1.2NA water immersion objective (488nm, 561nm and 640nm laser lines were used) and an Andor iXon+ EMCCD camera for image acquisition, respectively. For imaging of NS cells, the same DMEM/F12 was complemented with proliferation supplement, growth factors and laminin, and cells were grown in same chamber slides. 100μM Apcin was used instead for NS cells. For ChIP-seq validation results, Brn2-eGFP NS cells were synchronized with 330nM colchicine for 10 hours. Staining of DNA was done with far-red dye (Lukinavičius et al. 2015) - SiR-Hoechst (1:1000; prolonged timelapse) or Hoechst (1:10000 from 10mg/ml stock; snapshot acquisition). For imaging of NS cells, higher exposure times (>500ms) were often required. Post-imaging analysis, including background subtraction, was manually performed on time-lapse or snapshot data using Fiji software, using DNA staining to threshold chromosome signal. Mitotic chromosome enrichment (MCE) was calculated by dividing the mean fluorescence intensities of TF enrichment at metaphase chromatin and whole cells, and then converting data to log2 for visualization, as previously described (Teves et al. 2016). Scatter plots display mean values of MCE of all cells of a given population, in which each dot represents a single quantified cell.

### Tetracysteine-tag labelling

Ascl1/E47-TC and Brn2-TC expression vectors were generated by PCR amplification using primers shown in Supplemental Table S5 and Ascl1/E47 (Castro et al. 2006) and Brn2 template plasmids (Sugitani et al. 2002). For live-imaging, P19 cells were transfected with Ascl1/E47-TC or Brn2-TC expression vectors and synchronized with 60 μM proTAME and 200 μM Apcin. For labelling, cells were washed once with Opti-MEM (Life Technologies, #11058-021) and incubated with 2 μM ReAsH (Cayman, #19767) for 30 min at 37°C. ReAsH was removed and cells were washed again with Opti-MEM. A minimum of two 400 μM BAL incubations were carried out for 15 min at 37°C, in order to reduce non-specific ReAsH staining, with Opti-MEM washes in between. BAL was added a third time if cells saw good survival up to that point. After the last BAL incubation, cells were left in live-imaging medium.

### ChIP-seq and bioinformatics

Cultures at 70% confluency were incubated with 330nM colchicine for 10h. Synchronized NS cells were then gently “shaken-off” in order to enrich for the mitotic population. Thereafter cells were handled in 1,5mL tubes as typically only around 10×10^6^ cells were obtained from seven T-150 flasks of confluent NS cells. The same amount of interphase cells was gathered in parallel by using accutase on adherent cultures. Purity of our mitotic sample was confirmed in a small sample cytospined (Simport, #720-1962) and immunostained with anti-pHH3 (Merck, #06-570, 1:500) and DAPI. Only mitotic samples with >90% purity were further processed. For chromatin extraction from synchronized and unsynchronized cultures, cells were fixed sequentially with 2mM di(N-succinimidyl) glutarate (Sigma-Aldrich, #80424) and 1% formaldehyde (Sigma-Aldrich, #F8775) in phosphate-buffered saline and lysed, sonicated and immunoprecipitated as described (Castro et al. 2011), using rabbit anti-Brn2 antibody (Santa Cruz, #C-20). Library preparation (from precipitated material and input chromatin as control) was performed using NEBNext Ultra II DNA library Prep Kit (New England Biolabs, #E7645S) without size-selection to maintain sample complexity and according to manufacturer’s protocol. Next-generation sequencing was performed at Instituto Gulbenkian de Ciência (IGC) genomics facility using an Illumina NextSeq500 platform.

Dana analysis was performed in Galaxy (https://usegalaxy.eu/). Alignment against mm9 reference genome was done with Bowtie2, format conversion and removal of duplicate reads using SAMTools, and peak-calling using MACS 2.0 (Feng et al. 2012) with P value cut-off at 10(−10) using input chromatin as control. Subsampling of the data sets confirmed that peak calling saturation was achieved with approximately 80% of sequencing reads. Data visualization was done with the resulting bigwig files at the UCSC genome browser (http://genome.ucsc.edu/). Density plots and intersection of data sets were done using predefined tools available in Galaxy. *De novo* DNA motif search was performed using CisFinder with default settings (Sharov and Ko 2009) and Gene Ontology analysis using GREAT (McLean et al. 2010).

### Imaging and DNA colocalization analysis

eGFP fusion constructs were imaged at z-series stacks of 0.2μm, acquired at 37°C and 5% CO2, on a Yokogawa CSU-X Spinning Disk confocal, using a 60x 1.2NA water immersion objective. An Andor iXon+ EMCCD camera was used to acquire images for the emission of the eGFP (488 nm laser line). The colocalization of transcription factors with different DNA regions was analyzed with an image segmentation pipeline in FIJI. Cell nuclei were identified and segmented by k-means clustering based on the Hoechst signal. Three regions with high, medium and low Hoechst levels within each nucleus were defined, respectfully representing Heterochromatic, DNA dense, and DNA poor regions. Subsequently, the corresponding eGFP signal intensity in each of the 3 segmented regions was measured. Data is represented in log2 values, in which the intensity of GFP in each DNA region was divided by the whole cell intensity. At least twenty cells were analyzed per condition. Due to the quality of the Hoechst staining, DNA segmentation by K means clustering resulted in some variability among cells. Thus, only cells yielding a pre-defined segmentation ratio among the three different regions, Heterochromatin (10-20%), DNA dense (40-60%), and DNA poor (40%) were included in the final analysis. A max projection of the 20-30 0.2μm Z slices that were acquired, was applied before DNA segmentation.

### Fluorescence recovery after photobleaching (FRAP)

P19 cells grown and transfected as described above, were imaged at 37 °C and 5% CO2 using a Roper TIRF spinning disk confocal microscope with a 60×/1.27 NA water immersion objective, 488nm laser at 5% laser power, with 100–200 gain. Images were acquired at 250 × 250 pixels, with a pixel size of 0.21 μm. To image fluorescence recovery after photobleaching, a circular region of interest (ROI) with a diameter of 10 pixels was selected for bleaching, with 10 iterations (70 ms) of high intensity laser (50%). At least 9 pre-bleach fluorescence intensity values were averaged to normalize the post-bleach fluorescence recovery curve. Fluorescence recovery was then imaged for 25s or 50s at intervals of 0.50s. The recovery curve of the bleached ROI was normalized based on the intensity values before bleaching and on two more control ROIs: one for the background fluorescence intensity and the other for the fluorescent intensity of non-bleached cells. The t1/2 recovery time and mobile fraction were calculated using easyFRAP2mac (Rapsomaniki et al. 2012) and averaged over approximately 10 cells.

### Immunohistochemistry in embryo sections

E12.5 mouse heads were dissected and fixed in 4% PFA (Thermo fisher, #28906) for 60min at 4C°, washed once in PBS and dehydrated in 30% sucrose/PBS (Sigma-Aldrich, #S9378) at 4C° overnight. Tissue was then embedded in OCT (VWR #361603E) using a mold (Polysciences, #07918986), frozen and cut into cryosections. Sections of 12μm were obtained using a Leica cryostat CM-305S, collected on SuperFrost microscope slides (VWR, #631-0108) and stored at −20C°. For immunohistochemistry, slides were air-dried and then rehydrated in PBS for 5min at RT. Slides were blocked in PBS with 0,1% Triton X-100 (Sigma-Aldrich, #T8787) and 10% normal goat serum (Invitrogen, #10000C) for 30min at RT and then incubated with either anti-pHH3 (Merck, #06-570, 1:500) or anti-Ascl1 (Abcam, #ab211327, 1:500) and anti-βIII-Tubulin (Merck, #MAB1637, 1:200) in blocking solution overnight at 4C°. The following day, sections were washed 3×15min in PBS 0,1% Triton X-100 at RT and incubated with secondary antibodies in blocking solution for 2h at RT. Sections were washed 3×15min in PBS 0,01% Triton X-100 (PBST) and nuclei stained with DAPI (Sigma-Aldrich, #D9542, 1:10000) for 15min at RT after which they were washed once for 15min in PBST and 3×15min in PBS before mounting with Aqua-Poly/Mount (Polysciences, #07918606-20). All experiments were performed with wild-type mice from the NMRI strain and carried out upon approval and following the guidelines of the ethics committee of IGC.

### smRNA-FISH

Custom Stellaris FISH Probes were designed against Nestin and Dll1 by utilizing the Stellaris RNA FISH Probe Designer (biosearchtech.com/stellarisdesigner). Probes were coupled with either FAM, Q570 or Q670 fluorescent reporters (see Supplemental Table S6). RNA FISH protocol followed manufacturer’s instructions (biosearch.com/stellarisprotocols). Briefly, NS cells were grown on poly-L lysine (Sigma, #P8920) treated coverslips with thickness #0 (VWR, #631.0148). When confluency was reached, cells were fixed using RNase-free PBS-diluted 3,7% Formaldehyde (Sigma, #F8775), permeabilized with Triton 0,1% for 5min. and immunostained with anti-Ascl1 (Abcam, #ab211327) at 1:500 dilution (1h at RT). At this point, if RNase treatment was intended, permeabilized fixed NS cells were incubated for 20 minutes with RNase A (Invitrogen, #12091-021). NS cells were washed with designated buffers, and coverslips were incubated with the desired custom-made RNA FISH probes (at 1:100 concentration) during 16 hours at 37C° in the dark in a humidified chamber. In the next day, cells were washed, stained with DAPI and mounted in Vectashield Mounting Medium (Vector, #H-1000). Imaging was performed on a Deltavision widefield microscope coupled with a high sensitivity EM-CCD camera, using a 100x oil objective (1.4NA). Images acquired with Deltavision were deconvolved using Huygens Professional version 19.04 (Scientific Volume Imaging, The Netherlands, http://svi.nl) using standard parameters. In control experiments with transcriptional inhibitor, NS cells were incubated for 1 hour with 5μM triptolide added to fresh media prior to fixation. For DN-Brn2 exon counting, images were acquired using a Leica TCS SP8 confocal microscope equipped with two hybrid detectors for higher sensitivity and required lasers. A 63x (1.4 N.A.) oil objective was used, coupled with a zoom factor of 3. Images were analyzed using Fiji and exon and intron signal counting was performed using find maxima tool, defining the same threshold for all images. A single Z plane was used to confirm colocalization events and max projection of 25 Z stacks (0.3μm step size) used for exon/intron counting.

### Transcriptional and differentiation assays

Transcriptional assays in transfected P19 cells were performed using plasmids and protocols previously described (Castro et al. 2006). For differentiation assay, P19 cells were transfected with lipofectamine with expression vectors in chamber slides, and immunocytochemistry performed three days later as previously described (Vasconcelos et al. 2016).

## Accession numbers

ChIP-seq data sets reported in this study can be accessed under the code E-MTAB-9758 in Array Express.

## Author contributions

MAFS, Conception and experimental design, Data acquisition, Data analysis and interpretation, Drafting of manuscript; DSS, Conception and design, Acquisition of data, Data analysis and interpretation; VT, Data acquisition, Data analysis and interpretation; RBB and SMP, Contributed unpublished essential reagents and edited the manuscript; RAO, Conception and experimental design, Supervision, Drafting of manuscript; DSC, Conception and experimental design, Supervision, Drafting of manuscript.

## Acknowledgements

We thank Michael Wegner for providing Oct6-GFP, DBD (Oct6)-eGFP and DBD (Brn4)-eGFP plasmids. We thank Nuno Pimpão and Gabriel Martins at the IGC Advanced Imaging facility for excellent assistance with imaging acquisition and data analysis, the IGC Genomics facility for guidance with library synthesis and NGS, the IGC Flow Cytometry facility and Maria Azevedo at i3S Advanced Light Microscopy scientific platform. This work was funded by FCT grant PTDC/BIA-BID/29663/2017 to DSC and an FCT doctoral fellowship PD/BD/105999/2014 to MAFS.

## References

Abramo, Kristin, Anne-Laure Valton, Sergey V. Venev, Hakan Ozadam, A. Nicole Fox, and Job Dekker. 2019. “A Chromosome Folding Intermediate at the Condensin-to-Cohesin Transition during Telophase.” Nature Cell Biology 21 (11): 1393–1402.

Bertrand, Nicolas, Diogo S. Castro, and François Guillemot. 2002. “Proneural Genes and the Specification of Neural Cell Types.” Nature Reviews Neuroscience. https://doi.org/10.1038/nrn874.

Bylund, Magdalena, Elisabeth Andersson, Bennett G. Novitch, and Jonas Muhr. 2003. “Vertebrate Neurogenesis Is Counteracted by Sox1–3 Activity.” Nature Neuroscience. https://doi.org/10.1038/nn1131.

Cai, Yin, M. Julius Hossain, Jean-Karim Hériché, Antonio Z. Politi, Nike Walther, Birgit Koch, Malte Wachsmuth, et al. 2018. “Experimental and Computational Framework for a Dynamic Protein Atlas of Human Cell Division.” Nature 561 (7723): 411–15.

Caravaca, Juan Manuel, Greg Donahue, Justin S. Becker, Ximiao He, Charles Vinson, and Kenneth S. Zaret. 2013. “Bookmarking by Specific and Nonspecific Binding of FoxA1 Pioneer Factor to Mitotic Chromosomes.” Genes & Development 27 (3): 251–60.

Castro, Diogo S., Ben Martynoga, Carlos Parras, Vidya Ramesh, Emilie Pacary, Caroline Johnston, Daniela Drechsel, et al. 2011. “A Novel Function of the Proneural Factor Ascl1 in Progenitor Proliferation Identified by Genome-Wide Characterization of Its Targets.” Genes & Development 25 (9): 930–45.

Castro, Diogo S., Dorota Skowronska-Krawczyk, Olivier Armant, Ian J. Donaldson, Carlos Parras, Charles Hunt, James A. Critchley, et al. 2006. “Proneural bHLH and Brn Proteins Coregulate a Neurogenic Program through Cooperative Binding to a Conserved DNA Motif.” Developmental Cell 11 (6): 831–44.

Cong, Le, F. Ann Ran, David Cox, Shuailiang Lin, Robert Barretto, Naomi Habib, Patrick D. Hsu, et al. 2013. “Multiplex Genome Engineering Using CRISPR/Cas Systems.” Science 339 (6121): 819–23.

Conti, Luciano, Steven M. Pollard, Thorsten Gorba, Erika Reitano, Mauro Toselli, Gerardo Biella, Yirui Sun, et al. 2005. “Niche-Independent Symmetrical Self-Renewal of a Mammalian Tissue Stem Cell.” PLoS Biology 3 (9): e283.

Deluz, Cédric, Elias T. Friman, Daniel Strebinger, Alexander Benke, Mahé Raccaud, Andrea Callegari, Marion Leleu, Suliana Manley, and David M. Suter. 2016. “A Role for Mitotic Bookmarking of SOX2 in Pluripotency and Differentiation.” Genes & Development 30 (22): 2538–50.

Dugast-Darzacq, Claire, Sylvain Egloff, and Michel J. Weber. 2004. “Cooperative Dimerization of the POU Domain Protein Brn-2 on a New Motif Activates the Neuronal Promoter of the Human Aromatic L-Amino Acid Decarboxylase Gene.” Brain Research. Molecular Brain Research 120 (2): 151–63.

Feng, Jianxing, Tao Liu, Bo Qin, Yong Zhang, and Xiaole Shirley Liu. 2012. “Identifying ChIP-Seq Enrichment Using MACS.” Nature Protocols 7 (9): 1728–40.

Fernandez Garcia, Meilin, Cedric D. Moore, Katharine N. Schulz, Oscar Alberto, Greg Donague, Melissa M. Harrison, Heng Zhu, and Kenneth S. Zaret. 2019. “Structural Features of Transcription Factors Associating with Nucleosome Binding.” Molecular Cell 75 (5): 921–32.e6.

Festuccia, Nicola, Agnès Dubois, Sandrine Vandormael-Pournin, Elena Gallego Tejeda, Adrien Mouren, Sylvain Bessonnard, Florian Mueller, Caroline Proux, Michel Cohen-Tannoudji, and Pablo Navarro. 2016. “Mitotic Binding of Esrrb Marks Key Regulatory Regions of the Pluripotency Network.” Nature Cell Biology 18 (11): 1139–48.

Festuccia, Nicola, Inma Gonzalez, Nick Owens, and Pablo Navarro. 2017. “Mitotic Bookmarking in Development and Stem Cells.” Development 144 (20): 3633–45.

Festuccia, Nicola, Nick Owens, Thaleia Papadopoulou, Inma Gonzalez, Alexandra Tachtsidi, Sandrine Vandoermel-Pournin, Elena Gallego, et al. 2019. “Transcription Factor Activity and Nucleosome Organization in Mitosis.” Genome Research 29 (2): 250–60.

Finn, Elizabeth H., Gianluca Pegoraro, Hugo B. Brandão, Anne-Laure Valton, Marlies E. Oomen, Job Dekker, Leonid Mirny, and Tom Misteli. 2019. “Extensive Heterogeneity and Intrinsic Variation in Spatial Genome Organization.” Cell. https://doi.org/10.1016/j.cell.2019.01.020.

Friman, Elias T., Cédric Deluz, Antonio Ca Meireles-Filho, Subashika Govindan, Vincent Gardeux, Bart Deplancke, and David M. Suter. 2019. “Dynamic Regulation of Chromatin Accessibility by Pluripotency Transcription Factors across the Cell Cycle.” eLife 8 (December). https://doi.org/10.7554/eLife.50087.

Gebala, Magdalena, Stephanie L. Johnson, Geeta J. Narlikar, and Dan Herschlag. 2019. “Ion Counting Demonstrates a High Electrostatic Field Generated by the Nucleosome.” eLife 8 (June). https://doi.org/10.7554/eLife.44993.

Geoffroy, Cédric G., James A. Critchley, Diogo S. Castro, Sandra Ramelli, Christelle Barraclough, Patrick Descombes, Francois Guillemot, and Olivier Raineteau. 2009. “Engineering of Dominant Active Basic Helix-Loop-Helix Proteins That Are Resistant to Negative Regulation by Postnatal Central Nervous System Antineurogenic Cues.” Stem Cells 27 (4): 847–56.

Hsiung, Chris C-S, Caroline R. Bartman, Peng Huang, Paul Ginart, Aaron J. Stonestrom, Cheryl A. Keller, Carolyne Face, et al. 2016. “A Hyperactive Transcriptional State Marks Genome Reactivation at the Mitosis-G1 Transition.” Genes & Development 30 (12): 1423–39.

Imayoshi, Itaru, Akihiro Isomura, Yukiko Harima, Kyogo Kawaguchi, Hiroshi Kori, Hitoshi Miyachi, Takahiro Fujiwara, Fumiyoshi Ishidate, and Ryoichiro Kageyama. 2013. “Oscillatory Control of Factors Determining Multipotency and Fate in Mouse Neural Progenitors.” Science 342 (6163): 1203–8.

Jang, Hyonchol, Tae Wan Kim, Sungho Yoon, Soo-Youn Choi, Tae-Wook Kang, Seon-Young Kim, Yoo-Wook Kwon, Eun-Jung Cho, and Hong-Duk Youn. 2012. “O-GlcNAc Regulates Pluripotency and Reprogramming by Directly Acting on Core Components of the Pluripotency Network.” Cell Stem Cell 11 (1): 62–74.

Javasky, Elisheva, Inbal Shamir, Shashi Gandhi, Shawn Egri, Oded Sandler, Scott B. Rothbart, Noam Kaplan, Jacob D. Jaffe, Alon Goren, and Itamar Simon. 2018. “Study of Mitotic Chromatin Supports a Model of Bookmarking by Histone Modifications and Reveals Nucleosome Deposition Patterns.” Genome Research 28 (10): 1455–66.

Jerabek, Stepan, Calista Kl Ng, Guangming Wu, Marcos J. Arauzo-Bravo, Kee-Pyo Kim, Daniel Esch, Vikas Malik, et al. 2017. “Changing POU Dimerization Preferences Converts Oct6 into a Pluripotency Inducer.” EMBO Reports 18 (2): 319–33.

Josephson, R., T. Müller, J. Pickel, S. Okabe, K. Reynolds, P. A. Turner, A. Zimmer, and R. D. McKay. 1998. “POU Transcription Factors Control Expression of CNS Stem Cell-Specific Genes.” Development 125 (16): 3087–3100.

Kadauke, Stephan, Maheshi I. Udugama, Jan M. Pawlicki, Jordan C. Achtman, Deepti P. Jain, Yong Cheng, Ross C. Hardison, and Gerd A. Blobel. 2012. “Tissue-Specific Mitotic Bookmarking by Hematopoietic Transcription Factor GATA1.” Cell 150 (4): 725–37.

Kageyama, Ryoichiro, Toshiyuki Ohtsuka, Hiromi Shimojo, and Itaru Imayoshi. 2008. “Dynamic Notch Signaling in Neural Progenitor Cells and a Revised View of Lateral Inhibition.” Nature Neuroscience 11 (11): 1247–51.

Kalodimos, Charalampos G., Nikolaos Biris, Alexandre M. J. J. Bonvin, Marc M. Levandoski, Marc Guennuegues, Rolf Boelens, and Robert Kaptein. 2004. “Structure and Flexibility Adaptation in Nonspecific and Specific Protein-DNA Complexes.” Science 305 (5682): 386–89.

Kang, Hyeseon, Maxim N. Shokhirev, Zhichao Xu, Sahaana Chandran, Jesse R. Dixon, and Martin W. Hetzer. 2020. “Dynamic Regulation of Histone Modifications and Long-Range Chromosomal Interactions during Postmitotic Transcriptional Reactivation.” Genes & Development 34 (13-14): 913–30.

Liu, Yiyuan, Bobbie Pelham-Webb, Dafne Campigli Di Giammartino, Jiexi Li, Daleum Kim, Katsuhiro Kita, Nestor Saiz, et al. 2017. “Widespread Mitotic Bookmarking by Histone Marks and Transcription Factors in Pluripotent Stem Cells.” Cell Reports 19 (7): 1283–93.

Lodato, Michael A., Christopher W. Ng, Joseph A. Wamstad, Albert W. Cheng, Kevin K. Thai, Ernest Fraenkel, Rudolf Jaenisch, and Laurie A. Boyer. 2013. “SOX2 Co-Occupies Distal Enhancer Elements with Distinct POU Factors in ESCs and NPCs to Specify Cell State.” PLoS Genetics 9 (2): e1003288.

Lukinavičius, Gražvydas, Claudia Blaukopf, Elias Pershagen, Alberto Schena, Luc Reymond, Emmanuel Derivery, Marcos Gonzalez-Gaitan, et al. 2015. “SiR–Hoechst Is a Far-Red DNA Stain for Live-Cell Nanoscopy.” Nature Communications. https://doi.org/10.1038/ncomms9497.

Malik, Vikas, Dennis Zimmer, and Ralf Jauch. 2018. “Diversity among POU Transcription Factors in Chromatin Recognition and Cell Fate Reprogramming.” Cellular and Molecular Life Sciences: CMLS 75 (9): 1587–1612.

Martin, Brent R., Ben N. G. Giepmans, Stephen R. Adams, and Roger Y. Tsien. 2005. “Mammalian Cell-Based Optimization of the Biarsenical-Binding Tetracysteine Motif for Improved Fluorescence and Affinity.” Nature Biotechnology 23 (10): 1308–14.

McConnell, S. K., and C. E. Kaznowski. 1991. “Cell Cycle Dependence of Laminar Determination in Developing Neocortex.” Science 254 (5029): 282–85.

McEvilly, Robert J., Marcela Ortiz de Diaz, Marcus D. Schonemann, Farideh Hooshmand, and Michael G. Rosenfeld. 2002. “Transcriptional Regulation of Cortical Neuron Migration by POU Domain Factors.” Science 295 (5559): 1528–32.

McLean, Cory Y., Dave Bristor, Michael Hiller, Shoa L. Clarke, Bruce T. Schaar, Craig B. Lowe, Aaron M. Wenger, and Gill Bejerano. 2010. “GREAT Improves Functional Interpretation of Cis-Regulatory Regions.” Nature Biotechnology 28 (5): 495–501.

Miyagi, Satoru, Masazumi Nishimoto, Tetsuichiro Saito, Mikiko Ninomiya, Kazunobu Sawamoto, Hideyuki Okano, Masami Muramatsu, Hideyuki Oguro, Atsushi Iwama, and Akihiko Okuda. 2006. “The Sox2 Regulatory Region 2 Functions as a Neural Stem Cell-Specific Enhancer in the Telencephalon.” The Journal of Biological Chemistry 281 (19): 13374–81.

Nieto, Laurence, Gérard Joseph, Alexandre Stella, Pauline Henri, Odile Burlet-Schiltz, Bernard Monsarrat, Eric Clottes, and Monique Erard. 2007. “Differential Effects of Phosphorylation on DNA Binding Properties of N Oct-3 Are Dictated by protein/DNA Complex Structures.” Journal of Molecular Biology 370 (4): 687–700.

Owens, Nick, Thaleia Papadopoulou, Nicola Festuccia, Alexandra Tachtsidi, Inma Gonzalez, Agnes Dubois, Sandrine Vandormael-Pournin, et al. 2019. “CTCF Confers Local Nucleosome Resiliency after DNA Replication and during Mitosis.” eLife 8 (October). https://doi.org/10.7554/eLife.47898.

Palozola, Katherine C., Greg Donahue, Hong Liu, Gregory R. Grant, Justin S. Becker, Allison Cote, Hongtao Yu, Arjun Raj, and Kenneth S. Zaret. 2017. “Mitotic Transcription and Waves of Gene Reactivation during Mitotic Exit.” Science 358 (6359): 119–22.

Palozola, Katherine C., Jonathan Lerner, and Kenneth S. Zaret. 2019. “A Changing Paradigm of Transcriptional Memory Propagation through Mitosis.” Nature Reviews. Molecular Cell Biology 20 (1): 55–64.

Pilz, Gregor-Alexander, Atsunori Shitamukai, Isabel Reillo, Emilie Pacary, Julia Schwausch, Ronny Stahl, Jovica Ninkovic, et al. 2013. “Amplification of Progenitors in the Mammalian Telencephalon Includes a New Radial Glial Cell Type.” Nature Communications 4: 2125.

Raccaud, Mahé, Elias T. Friman, Andrea B. Alber, Harsha Agarwal, Cédric Deluz, Timo Kuhn, J. Christof M. Gebhardt, and David M. Suter. 2019. “Mitotic Chromosome Binding Predicts Transcription Factor Properties in Interphase.” Nature Communications 10 (1): 487.

Raccaud, Mahé, and David M. Suter. 2018. “Transcription Factor Retention on Mitotic Chromosomes: Regulatory Mechanisms and Impact on Cell Fate Decisions.” FEBS Letters 592 (6): 878–87.

Raposo, Alexandre A. S. F., Francisca F. Vasconcelos, Daniela Drechsel, Corentine Marie, Caroline Johnston, Dirk Dolle, Angela Bithell, et al. 2015. “Ascl1 Coordinately Regulates Gene Expression and the Chromatin Landscape during Neurogenesis.” Cell Reports 10 (9): 1544–56.

Rapsomaniki, Maria Anna, Panagiotis Kotsantis, Ioanna-Eleni Symeonidou, Nickolaos-Nikiforos Giakoumakis, Stavros Taraviras, and Zoi Lygerou. 2012. “easyFRAP: An Interactive, Easy-to-Use Tool for Qualitative and Quantitative Analysis of FRAP Data.” Bioinformatics 28 (13): 1800–1801.

Saxe, Jonathan P., Alexey Tomilin, Hans R. Schöler, Kathrin Plath, and Jing Huang. 2009. “Post-Translational Regulation of Oct4 Transcriptional Activity.” PloS One 4 (2): e4467.

Sharov, Alexei A., and Minoru S. H. Ko. 2009. “Exhaustive Search for over-Represented DNA Sequence Motifs with CisFinder.” DNA Research: An International Journal for Rapid Publication of Reports on Genes and Genomes 16 (5): 261–73.

Shimojo, Hiromi, Akihiro Isomura, Toshiyuki Ohtsuka, Hiroshi Kori, Hitoshi Miyachi, and Ryoichiro Kageyama. 2016. “Oscillatory Control of Delta-like1 in Cell Interactions Regulates Dynamic Gene Expression and Tissue Morphogenesis.” Genes & Development 30 (1): 102–16.

Shin, Chanseok, and James L. Manley. 2002. “The SR Protein SRp38 Represses Splicing in M Phase Cells.” Cell. https://doi.org/10.1016/s0092-8674(02)01038-3.

Shin, Jihoon, Tae Wan Kim, Hyunsoo Kim, Hye Ji Kim, Min Young Suh, Sangho Lee, Han-Teo Lee, et al. 2016. “Aurkb/PP1-Mediated Resetting of Oct4 during the Cell Cycle Determines the Identity of Embryonic Stem Cells.” eLife 5 (February): e10877.

Soufi, Abdenour, and Stephen Dalton. 2016. “Cycling through Developmental Decisions: How Cell Cycle Dynamics Control Pluripotency, Differentiation and Reprogramming.” Development 143 (23): 4301–11.

Steffen, Philipp A., and Leonie Ringrose. 2014. “What Are Memories Made of? How Polycomb and Trithorax Proteins Mediate Epigenetic Memory.” Nature Reviews. Molecular Cell Biology 15 (5): 340–56.

Sugitani, Yoshinobu, Shigeyasu Nakai, Osamu Minowa, Miyuki Nishi, Kou-Ichi Jishage, Hitoshi Kawano, Kensaku Mori, Masaharu Ogawa, and Tetsuo Noda. 2002. “Brn-1 and Brn-2 Share Crucial Roles in the Production and Positioning of Mouse Neocortical Neurons.” Genes & Development 16 (14): 1760–65.

Sunabori, Takehiko, Akinori Tokunaga, Takeharu Nagai, Kazunobu Sawamoto, Masaru Okabe, Atsushi Miyawaki, Yumi Matsuzaki, Takaki Miyata, and Hideyuki Okano. 2008. “Cell-Cycle-Specific Nestin Expression Coordinates with Morphological Changes in Embryonic Cortical Neural Progenitors.” Journal of Cell Science 121 (Pt 8): 1204–12.

Suter, David M. 2020. “Transcription Factors and DNA Play Hide and Seek.” Trends in Cell Biology 30 (6): 491–500.

Tanaka, Shinya, Yusuke Kamachi, Aki Tanouchi, Hiroshi Hamada, Naihe Jing, and Hisato Kondoh. 2004. “Interplay of SOX and POU Factors in Regulation of the Nestin Gene in Neural Primordial Cells.” Molecular and Cellular Biology 24 (20): 8834–46.

Teves, Sheila S., Luye An, Anders S. Hansen, Liangqi Xie, Xavier Darzacq, and Robert Tjian. 2016. “A Dynamic Mode of Mitotic Bookmarking by Transcription Factors.” eLife 5 (November). https://doi.org/10.7554/eLife.22280.

Vasconcelos, Francisca F., and Diogo S. Castro. 2014. “Transcriptional Control of Vertebrate Neurogenesis by the Proneural Factor Ascl1.” Frontiers in Cellular Neuroscience. https://doi.org/10.3389/fncel.2014.00412.

Vasconcelos, Francisca F., Alessandro Sessa, Cátia Laranjeira, Alexandre A. S. F. Raposo, Vera Teixeira, Daniel W. Hagey, Diogo M. Tomaz, Jonas Muhr, Vania Broccoli, and Diogo S. Castro. 2016. “MyT1 Counteracts the Neural Progenitor Program to Promote Vertebrate Neurogenesis.” Cell Reports 17 (2): 469–83.

Wapinski, Orly L., Thomas Vierbuchen, Kun Qu, Qian Yi Lee, Soham Chanda, Daniel R. Fuentes, Paul G. Giresi, et al. 2013. “Hierarchical Mechanisms for Direct Reprogramming of Fibroblasts to Neurons.” Cell 155 (3): 621–35.

Zaret, Kenneth S. 2014. “Genome Reactivation after the Silence in Mitosis: Recapitulating Mechanisms of Development?” Developmental Cell 29 (2): 132–34.

Zhang, Haoyue, Daniel J. Emerson, Thomas G. Gilgenast, Katelyn R. Titus, Yemin Lan, Peng Huang, Di Zhang, et al. 2019. “Chromatin Structure Dynamics during the Mitosis-to-G1 Phase Transition.” Nature 576 (7785): 158–62.

