## Supplemental Figures and Tables for "Hierarchical reactivation of transcription during mitosis-to-G1 transition by Brn2 and Ascl1 in neural stem cells"

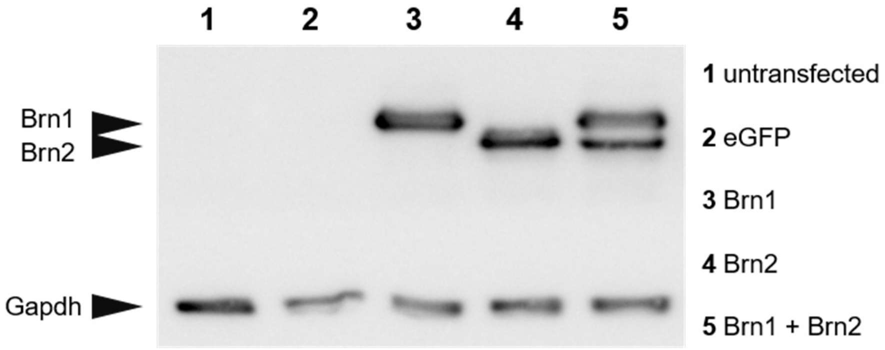


**Supplemental Figure S1. Commercial antibody against Brn2 also recognizes Brn1.** P19 cells were either not transfected or transfected with expression plasmids for eGFP, Brn1, Brn2 or Brn1 and Brn2, as indicated in the figure. Western-blot was performed with commercial antibody used in Figure 1E and Supplemental Figure S6A. Gapdh was used as loading control. Black triangles mark the position of Brn1, Brn2 and Gapdh specific bands, as indicated in the figure.

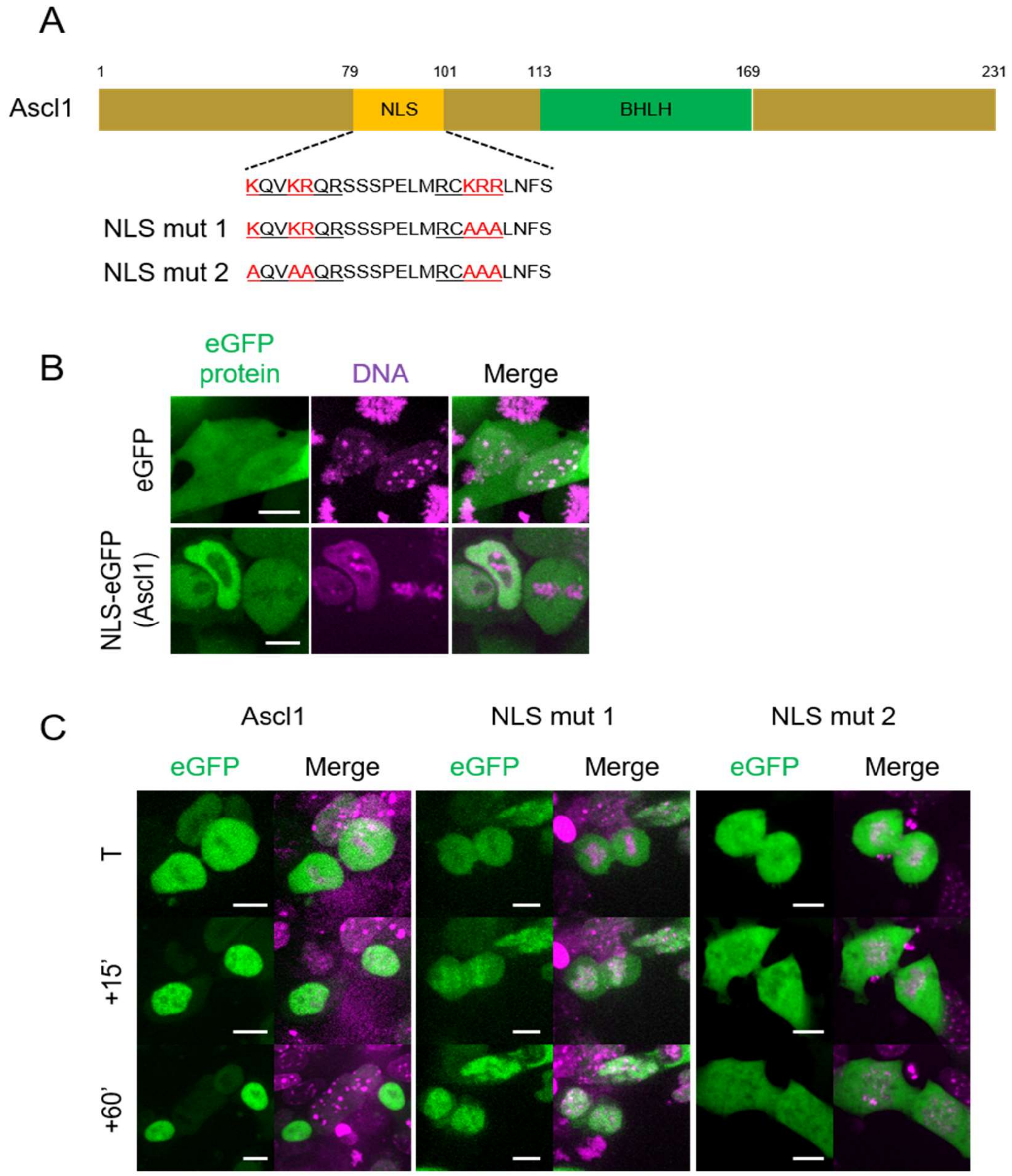


**Supplemental Figure S2. Colocalization of Ascl1 with chromatin at late telophase/early G1 requires active nuclear import. (A)** Cartoon depicting the sequence of the identified Ascl1 bipartite NLS (using nls-mapper.iab.keio.ac.jp), and mutations inserted when generating Ascl1 NLS mutants. **(B)** Live-cell imaging analysis of P19 cells transfected with eGFP or NLS(Ascl1)-eGFP, as indicated in figure and in presence of DNA dye SiR-Hoechst **(C)** Time-lapse from live-cell imaging analysis of P19 cells transfected with full-length Ascl1 (or Ascl1 NLS mutants as described above) in fusion with eGFP, as indicated in figure and in presence of DNA dye SiR-Hoechst. T= Telophase. Scale bar=10µm.

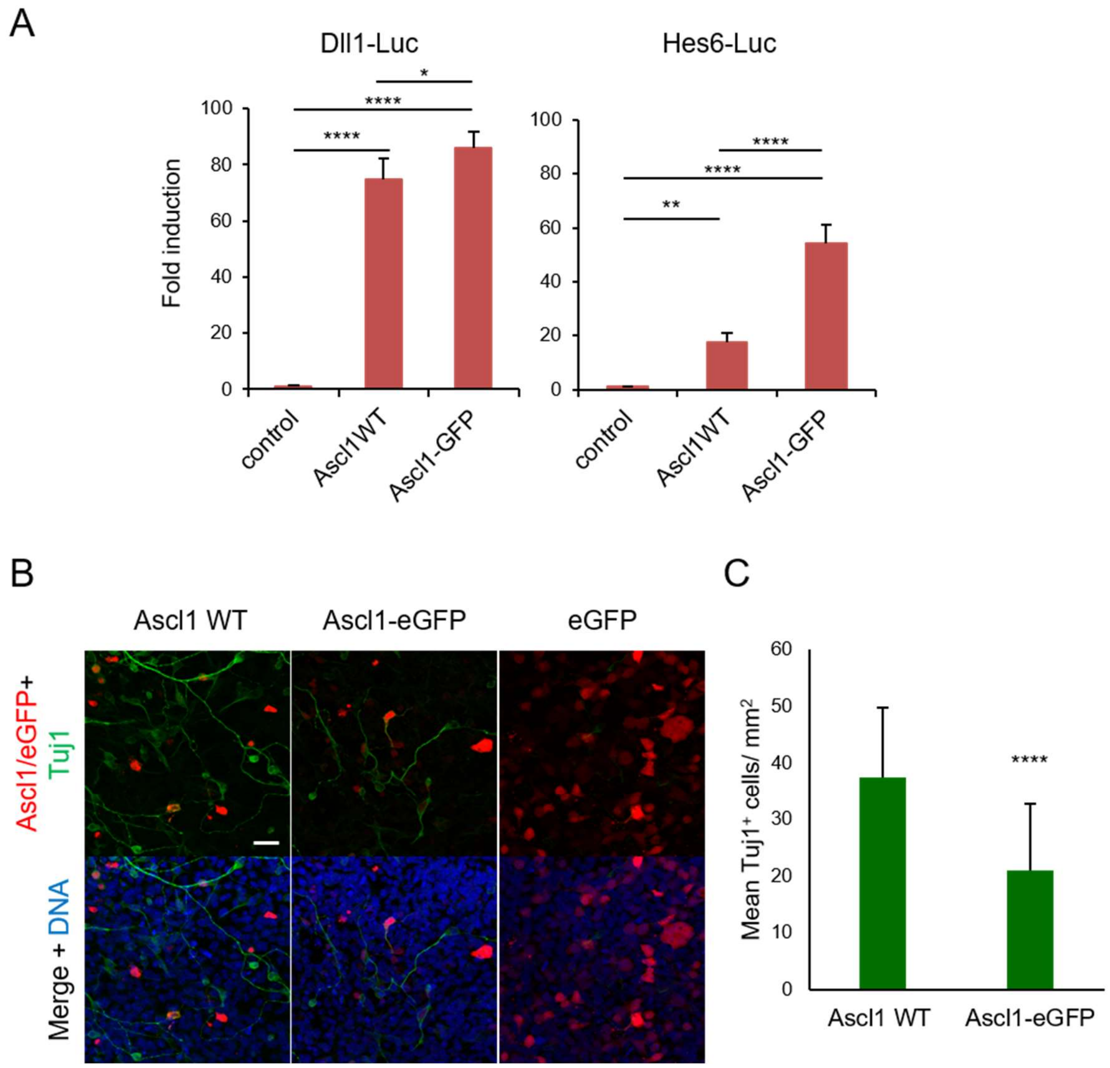


**Supplemental Figure S3. Ascl1-eGFP fusion protein is functional. (A)** Transcriptional assays performed in P19 cells co-transfected with vectors containing luciferase reporter gene under the regulation of regulatory regions from Dll1 (Left) and Hes6 (Right) genes, and Ascl1 or Ascl1-eGFP expression plasmids. Data are shown as mean± SD of four biological replicates. **(B)** Representative images from immunocytochemistry analysis for expression of neuronal marker Tuj1 in P19 cells 72 hours upon transfection with Ascl1-IRES-GFP, Ascl1-eGFP or eGFP expression plasmids. **(C)** Mean number of immature neurons (Tuj1$^+$) per mm$^2$ in both Ascl1 WT and Ascl1-eGFP after subtraction of mean number of Tuj1$^+$ cells found in the control (eGFP). n=32 images for each condition. One-way ANOVA Tukey's multiple comparison test (A) and Mann-Whitney test (C) were performed. $p>0.05$ (n.s.), $p\leq0.05$ (*), $p\leq0.01$ (**), $p\leq0.001$ (***), $p\leq0.0001$ (****). Scale bar=30µm.

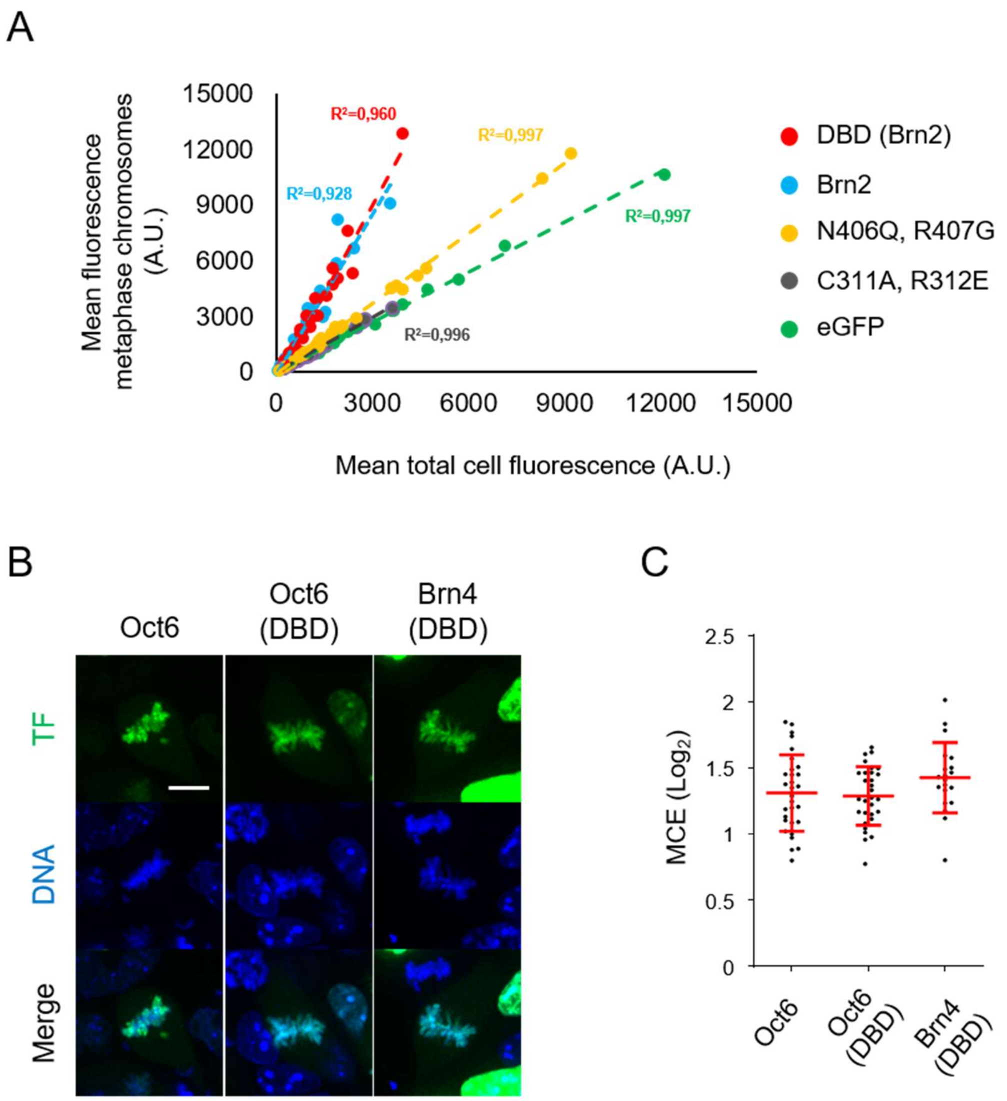


**Supplemental Figure S4. (A) Mitotic chromosome enrichment increases linearly with expression levels, in transfected P19 cells.** Total cell mean fluorescence was plotted as a function of metaphase mean fluorescence, using raw values used in MCE quantifications shown in Figure 4. The correlation coefficient (R) associated with each Brn2 protein suggests linear variation. **(B-C) The DNA binding domain of POU3Fs is sufficient for mitotic chromosome binding**. **(B)** Representative captures from live-cell imaging analysis of P19 cells expressing Oct6 or truncated DBDs of Oct6 and Brn4, in fusion with eGFP. Imaging was performed upon synchronization of cells using proTAME and Apcin and in the presence of DNA staining Hoechst. **(C)** Quantifications of mitotic chromosome enrichment levels shown in (B). Data shown as mean ± SD (n=30 cells for Oct6 full length and DBD; n=20 for Brn4 DBD). Scale bar=10µm.

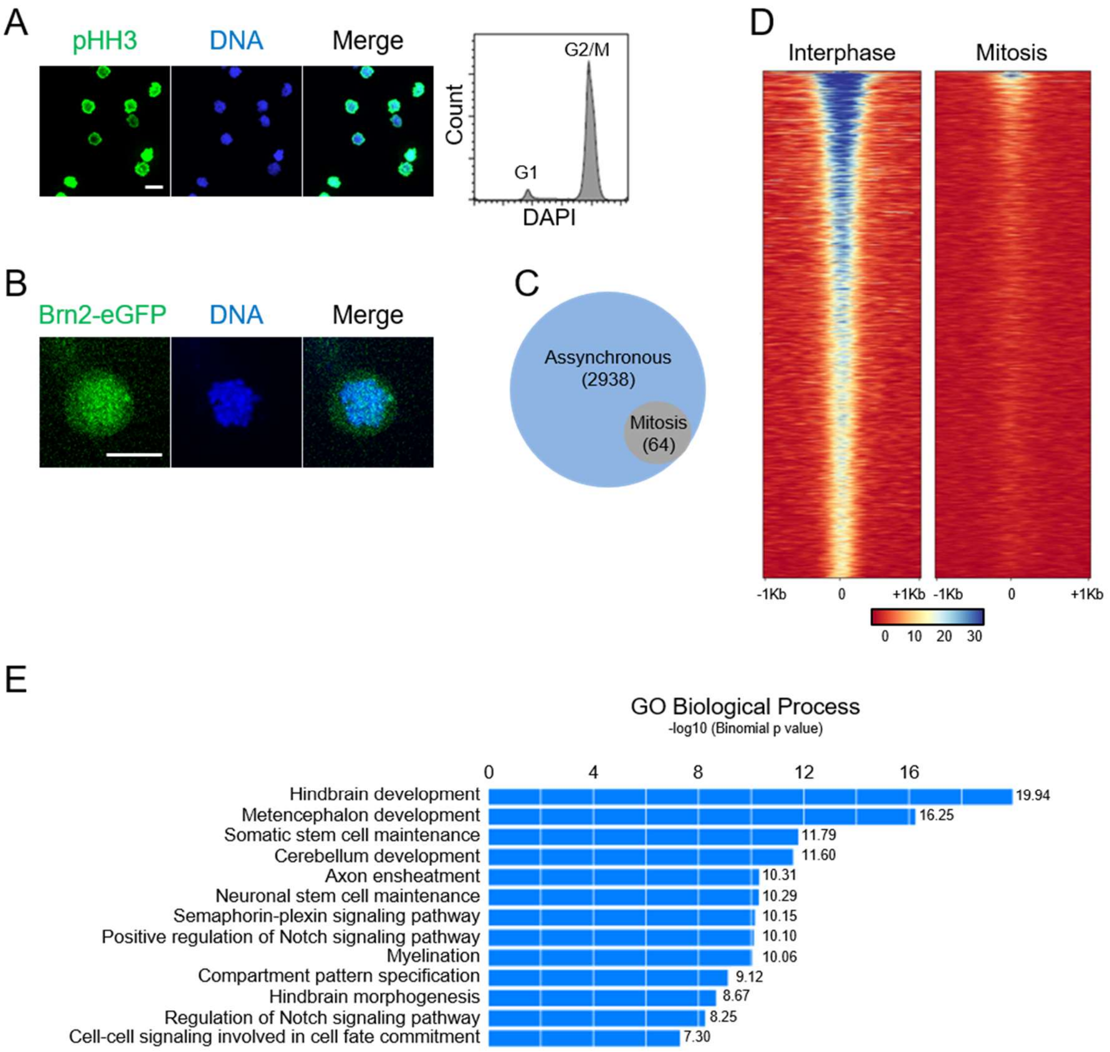


**Supplemental Figure S5. Genome-wide profiling of Brn2 binding in interphase and mitotic chromatin using ChIP-seq. (A)** Co-staining for phospho-histone H3 and DAPI (left) and FACS analysis (right) of NS cells upon a synchronization protocol that combines colchicine with shake-off, resulting consistently in a mitotic fraction of 90-95%. **(B)** Representative image showing association of eGFP-tagged Brn2 with mitotic chromatin is maintained upon synchronization of NS cells with colchicine. **(C)** Venn diagram depicting number of Brn2 peaks found by ChIP-seq in mitotically arrested or asynchronous NS cultures, in a replicate experiment to the one described in Figure 5, using a 90% mitotically enriched cell population. **(D)** Density plot of ChIP-seq reads from mitotic and interphase samples from second replicate experiment, mapping to genomic regions centered on Brn2 peak summits found in asynchronous sample (see Figure 5 legend for details). **(E)** Gene ontology (GO) analysis run on list of Brn2 bound target genes in interphase, finds "somatic" and "neuronal stem cell maintenance" at the top of enriched biological process terms. Scale bars=10µm.

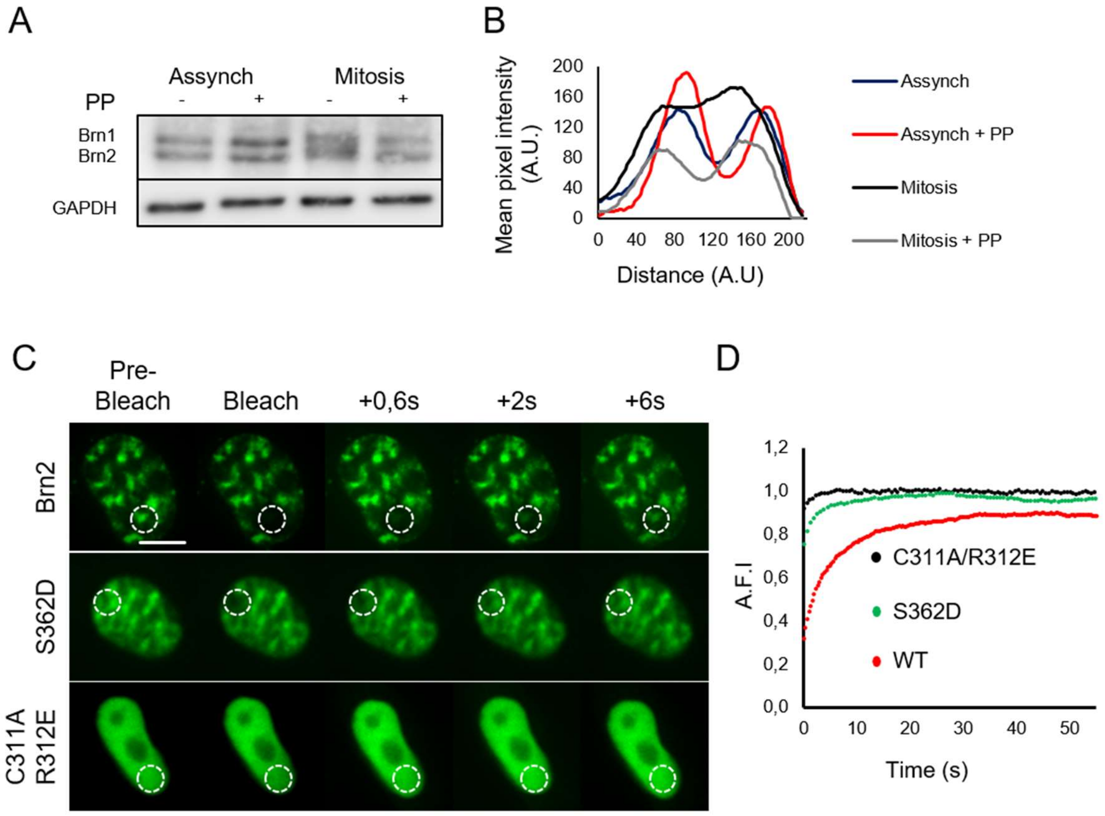


E

| | $T_{1/2}$ Recovery Time (sec) | Mobile Fraction (%) |
|---|---|---|
| WT Brn2 | 3.66 ± 1.6 | 0.85 ± 0.8 |
| C311A R312E | 0.67 ± 0.33 | 0.86 ± 0.19 |
| S362D | 1.71 ± 1.06 | 0.87 ± 0.32 |

**Supplemental Figure S6. (A-B) Analysis of phospho-status of Brn2 in NS cells**. **(A)** Western-blot with an antibody recognizing Brn1/2, using protein extracts from mitotically arrested or non-synchronized NS cultures, and with or without treatment with Lambda phosphatase (PP). Gapdh was used as a loading control. **(B)** Quantification of western-blot from C using mean pixel intensity reveals a strong smear in between Brn1 and Brn2 bands in mitotic extracts, which disappears upon PP treatment. **(C-E)** FRAP assay performed in P19 cells transfected with Brn2 derivatives. **(C)** P19 cells were transfected with expression plasmids for eGFP fusions of Brn2 or derivatives (C311A/R312E; S362A or S362D) and images taken before and after bleaching at timepoints indicated in figure. **(D-E)** Data was normalized and plotted in order to find the $t_{1/2}$ recovery time and mobile fraction for each Brn2 derivative. Data shown as mean± SD (n=10 except for Brn2 data that derives from 9 cells). See Materials and Methods for further details on data analysis.

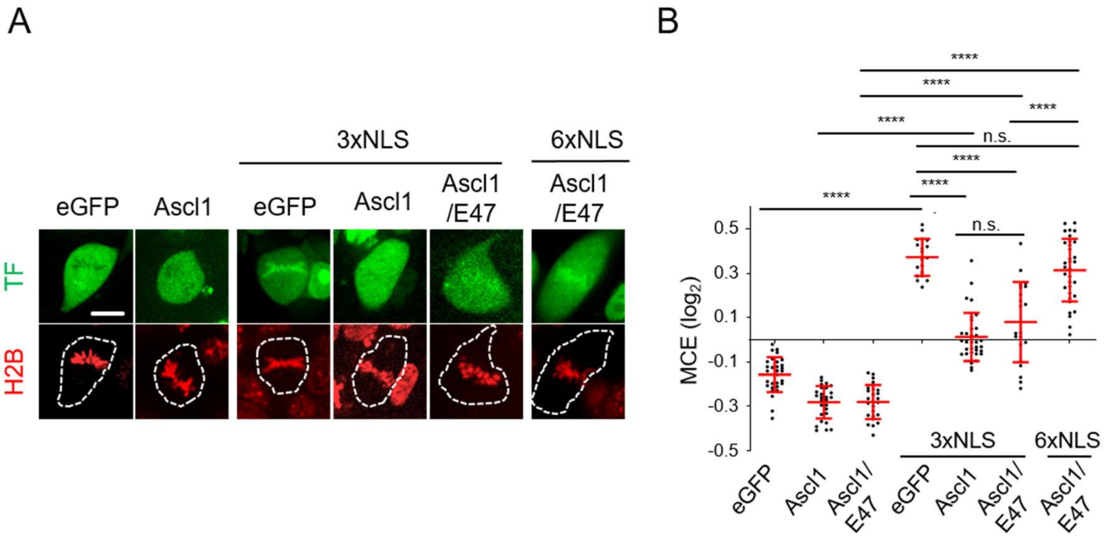


**Supplemental Figure S7. Live-cell imaging and DNA colocalization analysis of Ascl1 derivatives in fusion with SV40-NLS sequences. (A)** Representative captures from live-cell imaging of P19-H2B-RFP cells expressing eGFP, Ascl1 or derivatives fused to SV40 NLS sequences. Cells were synchronized in metaphase using proTAME and Apcin and live imaged. H2B-RFP was used to visualize chromatin. **(B)** Quantifications of mitotic chromosome enrichment levels for images taken in (A). N=30 (eGFP), 27 (Ascl1), 25 (Ascl1/E47), 19 (eGFP-3xNLS), 33 (Ascl1-3xNLS), 21 (Ascl1/E47-3xNLS) and 33 cells (Ascl1/E47-6xNLS). Scale bar=10µm.

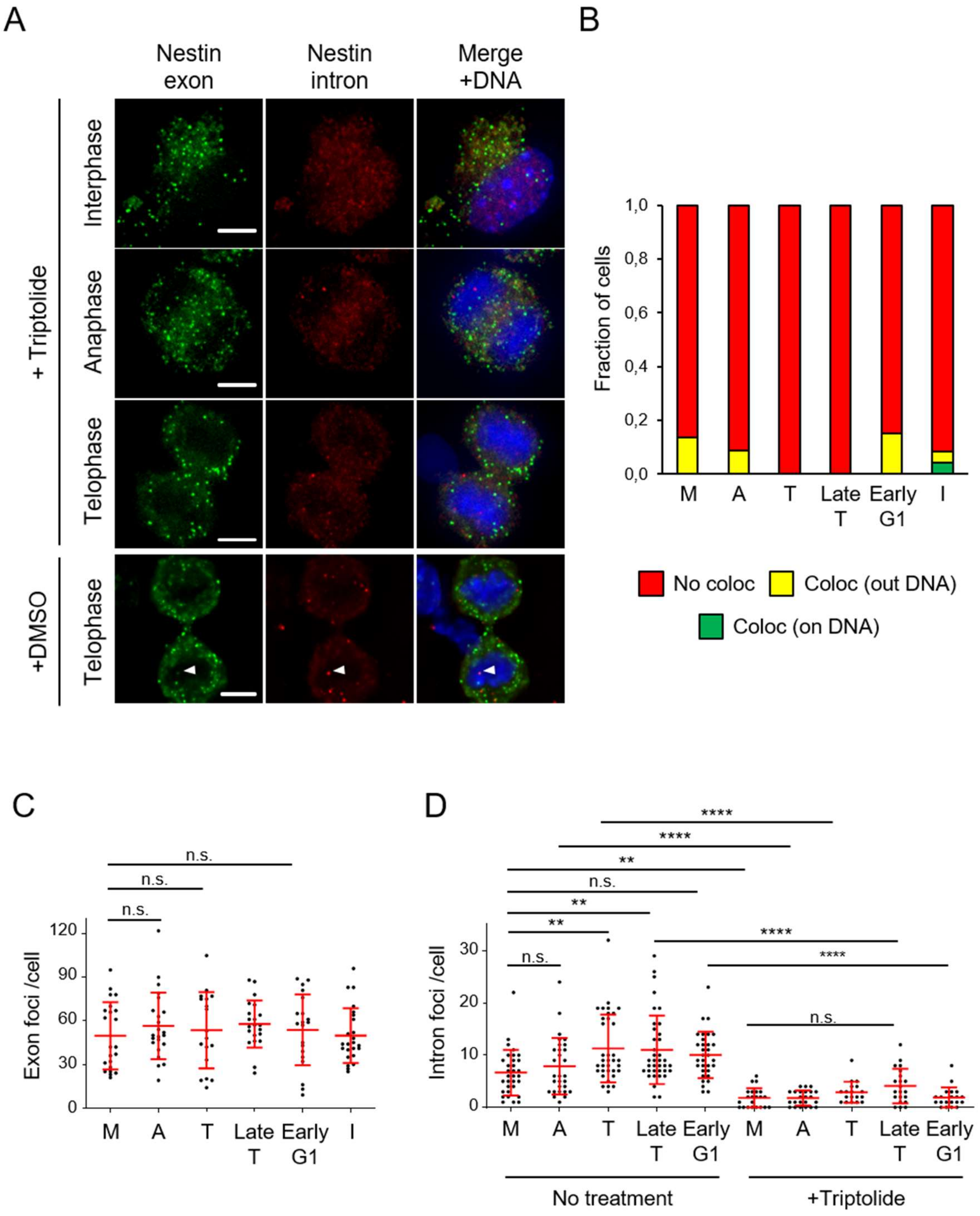


**Supplemental Figure S8. Incubation with the transcriptional inhibitor triptolide abolishes evidence of active-transcription in NS cells as detected by smRNA-FISH. (A)** Representative images of NS cells in interphase, anaphase and telophase, incubated with triptolide (1h) or vehicle control (DMSO) as indicated in figure, and stained by smRNA-FISH using exonic (FAM) and intronic (Q570) probes for Nestin transcript. DNA was stained with DAPI. Images shown are maximum intensity projections of six optical planes with 0,3μm Z step intervals. Scale bars=5µm. **(B)** Stacked bar plots showing the fractions of cells containing at least one spot of colocalized intron-exon probe signal found on DNA (green), outside DNA (yellow) or without colocalization spots (red) (n=22 in metaphase; n=23 in anaphase; n=18 in telophase, n=21 in late telophase; n=20 in early G1; n=24 in interphase). **(C)** Quantifications of exon probe signal per cell for Nestin transcripts at different stages during MG1 transition in cells incubated with triptolide and partially shown in (A). (D) Quantifications of intron probe signal per cell for Nestin transcripts at different stages during MG1 transition in cells incubated with triptolide or control conditions (same cells as represented in Figure 7) One-way ANOVA Tukey's multiple comparison test was performed. $p>0.05$ (n.s.), $p\leq0.05$ (*), $p\leq0.01$ (**), $p\leq0.001$ (***), $p\leq0.0001$ (****). M=metaphase; A=anaphase; T=telophase; Late T=late telophase; I=interphase. Data shown as mean ± SD.

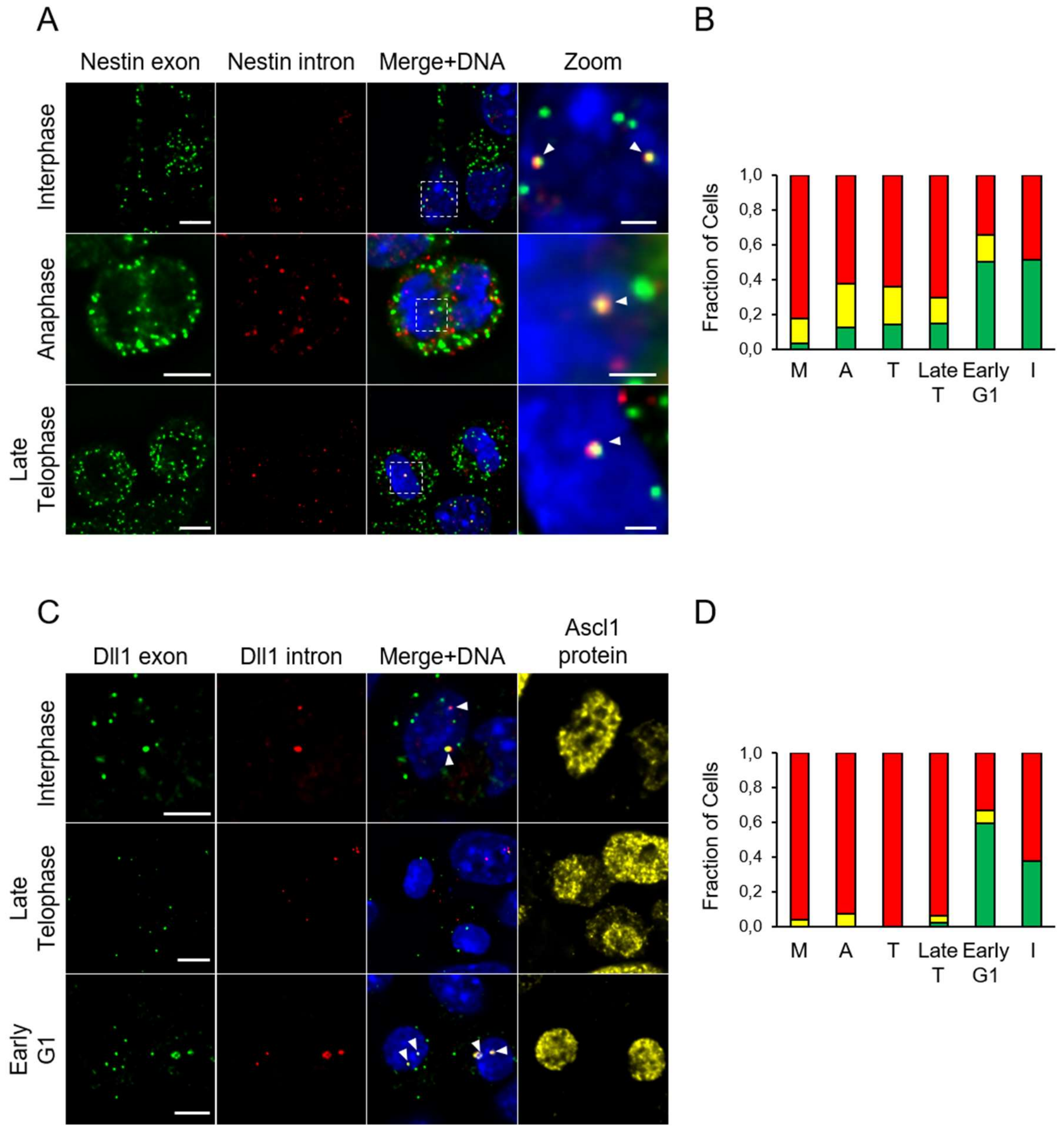


**Supplemental Figure S9. Validation of smRNA-FISH showing active transcription of Nestin and Dll1 genes in NS cells, using probes with different fluorophores. (A)** Representative images of cells in interphase, anaphase and late telophase stained by smRNA-FISH using exonic (Q570) and intronic (Q670) probes for Nestin transcript. Images shown are maximum intensity projections of six optical planes with 0,3µm Z step intervals, with white arrow-heads marking spots of colocalized intron-exon probe signal found on DNA (stained with DAPI) Scale bars=5µm. **(B)** Stacked bar plots showing the fractions of cells containing at least one spot of colocalized intron-exon probe signal found on DNA (green), outside DNA (yellow) or without colocalization spots (red) (n=62 in metaphase; n=56 in anaphase; n=42 in telophase, n=34 in late telophase; n=26 in early G1; n=66 in interphase). **(C)**

Representative images of cells in interphase, late telophase and early G1 co-stained by smRNA-FISH using exonic (Q670) and intronic (Q570) probes for Dll1 transcript, and by immuno-cytochemistry for Ascl1 protein. Images shown are maximum intensity projections of six optical planes with 0,3μm Z step intervals, with white arrow-heads marking spots of colocalized intron-exon probe signal found on DNA (stained with DAPI). Scale bars=5μm. **(D)** Stacked bar plots showing the fraction of cells containing at least one spot of colocalized intron/exon probe signal found on DNA (green), outside DNA (yellow) or without colocalization spots (red) (n=73 in metaphase; n=42 in anaphase; n=42 in telophase, n=48 in late telophase; n=27 in early G1; n=32 in interphase).

### Supplemental tables

#### Supplemental Table S1 – PCR primers used in subcloning

| Plasmids | Primers (5´-3´) | Restriction enzymes |
|---|---|---|
| **Ascl1-eGFP** | **Fw:** ATC ATC CTC GAG CGG CCC GGC ATG GAG AGC TCT<br>**Rev:** GAT GAT CCC GGG CGA ACC AGT TGG TAA AGT CCA GCA GCT CTT GTT CCT C | XhoI<br>XmaI |
| **Ascl1/E47-eGFP** | **Fw**: ATC ATC CTC GAG CGG CCC GGC ATG GAG AGC GGA GCC GGC C<br>**Rev**: GAT GAT ACC GGT CCC AGG TGC CCG GCT GGG TTG TG | XhoI<br>AgeI |
| **Brn2-eGFP** | **Fw:** ATC ATC GAA TTC GAG AGT C ATG GCG ACC GCA GCG TCT AAC CAC<br>**Rev:** GAT GAT GGA TCC CC CTG GAC GGG CGT CTG CAC CCC | EcoRI<br>BAMHI |
| **DBD (Brn2)-eGFP** | **Fw**: ATC ATC GAA TTC GAG AGT CAT GCC GGG CCA CCC AGG CGC GCA C<br>**Rev:**GAT GAT GGA TCC CCA CCC CCA TAC ACA TCC TCG GC | EcoRI<br>BAMHI |

#### Supplemental Table S2 – Primers used in site-directed mutagenesis

| Plasmids | Primers (5´-3´) |
|---|---|
| **NLS mutant 1-eGFP** | **Fw**: CCG GAA CTG ATG CGC TGC GCA GCC GCG CTC AAC TTC AGC GGC TTC GGC<br>**Rev:** GCC GAA GCC GCT GAA GTT GAG CGC GGC TGC GCA GCG CAT CAG TTC CGG |
| **NLS mutant 2-eGFP** | **Fw**: GTC ACA AGT CAG CGG CCG CGC AGG TCG CGG CCC AGC GCT CGT CCT C |

| | |
|---|---|
| | **Rev**: G AGG ACG AGC GCT GGG CCG CGA CCT GCG CGG CCG CTG ACT TGT GAC |
| **M414N-eGFP** | **Fw:** CAG AAA GAG AAA AGG AAT ACC CCT CCC GGA GGG<br>**Rev:** CCC TCC GGG AGG GGT ATT CCT TTT CTC TTT CTG |
| **S362A-eGFP** | **Fw:** AAG CGG ACC GCC ATC GAG GTG<br>**Rev:** CAC CTC GAT GGC GGT CCG CTT |
| **S362D-eGFP** | **Fw:** AAG CGG ACC GAC ATC GAG GTG AGC<br>**Rev:** GCT CAC CTC GAT GTC GGT CCG CTT |

#### Supplemental Table S3 – Oligonucleotide sequences used in NLS constructs

| | **Oligonucleotide sequences (5´-3´)** |
|---|---|
| **NLS (Ascl1)-eGFP** | **Fw:**GTACATTAAGCAGGTCAAGCGCCAGCGCTCGTCCTCTCCGGAACTGATGCGCTGCAAACGCCGGCTCAACTTCAGCTAACTCGAGT<br>**Rev:**CTAGACTCGAGTTAGCTGAAGTTGAGCCGGCGTTTGCAGCGCATCAGTTCCGGAGAGGACGAGCGCTGGCGCTTGACCTGCTTAAT |
| **3xSV40 NLS-eGFP** | **Fw**:GTACATTGATCCAAAAAAGAAGAGAAAGGTAGATCCAAAAAAGAAGAGAAAGGTAGATCCAAAAAAGAAGAGAAAGTGACTCGAGT<br>**Rev:**CTAGACTCGAGTCACTTTCTCTTCTTTTTTGGATCTACCTTTCTCTTCTTTTTTGGATCTACCTTTCTCTTCTTTTTTGGATCAAT |

#### Supplemental Table S4 – Primers used in Gibson assembly for generation of CRISPR/Cas9 vectors

| | |
|---|---|
| **Ascl1_LHA_Fw** | CGGCCGCTCTAGAACTAGTGGAAGGCGCCGGCGGC |
| **Ascl1_LHA_Rev** | CCTTGCTCACCATGAACCAGTTGGTAAAGTCCAGCAGCTCTTGTTCCTCTGGGC |
| **Ascl1_EGFP_Fw** | CCAACTGGTTCATGGTGAGCAAGGGCGAGGAGCTGTTCACCGGG |
| **Ascl1_EGFP_Rev** | CCTGGCAGGTCCTTACTTGTACAGCTCGTCCATGCCGAGAGTGATCCC |
| **Ascl1_RHA_Fw** | CTGTACAAGTAAGGACCTGCCAGGCTCTCCTGGGAATGGACTTTGGAAGCAGG |

| | |
|---|---|
| **Ascl1_RHA_Rev** | CCTCGAGGTCGACGGTATAAAAAATTCACAAAGATTGGAAGCATT A |
| **Brn2_LHA_Fw** | CGGCCGCTCTAGAACGACGAGCTGCACGGGCC |
| **Brn2_LHA_Rev** | CCTTGCTCACCATCTGGACGGGCGTCTGCACCCCGTGGTGTGG |
| **Brn2_EGFP_Fw** | ACGCCCGTCCAGATGGTGAGCAAGGGCGAGGAGCTGTTCACCG GG |
| **Brn2_EGFP_Rev** | GCCCCTCCCCCGCTTGAGTTTACTTGTACAGCTCGTCCATGCCGA GAGTGATCCCGGCGGC |
| **Brn2_RHA_Fw** | CTGTACAAGTAAACTCAAGCGGGGGAGGGGCAGAGCTTGGGGCT CC |

**Supplemental Table S5 - Primers used in generation of tetracysteine-tag fusion proteins**

| | |
|---|---|
| **TC-Brn2-IRES-GFP** | **Fw:** ATC ATC GAA TTC TGG CCC GGC ATG TGT TGC CCG GGC TGC TGT ATG GAG AGC TCT GGC AAG ATG GAG AGT GGA<br><br>**Rev:** GAT GAT CTC GAG TCA GAA CCA GTT GGT AAA GTC CAG CAG CTC TTG TTC |
| **TC-Ascl1-E47-IRES-GFP** | **Fw:** ATC ATC CTC GAG CGG CCC GGC ATG TTC CTG AAC TGC TGC CCG GGT TGC TGC AT GGA ACC GAT GGA GAG CGG AGC CGG CCA GCA G<br><br>**Rev:** GAT GAT GAT ATC TCA CAG GTG CCC GGC TGG GTT GTG |

**Supplemental Table S6 - Dll1 exon smRNA FISH probe sequences**

| | | | |
|---|---|---|---|
| tcagctcaaatacgccggag | Dll1 exon_1 | tagaacaaggggaagagccg | Dll1 exon_25 |
| ttcttgttgacgaactcctg | Dll1 exon_2 | cacaggtaagagttgccgag | Dll1 exon_26 |
| gaaagaaggtcctgcaggcg | Dll1 exon_3 | acagtcatccacattgtcct | Dll1 exon_27 |
| tggtagtgcttgaggcatac | Dll1 exon_4 | catttgcacacggggaggag | Dll1 exon_28 |
| caggctgaaggagtcgacac | Dll1 exon_5 | caggtacaggagaagtcgtt | Dll1 exon_29 |
| aatcggatggggttgctgaa | Dll1 exon_6 | cactcacacatgtagcgctg | Dll1 exon_30 |
| attcttctcccacagtgagg | Dll1 exon_7 | gagcagaaactggcagttgg | Dll1 exon_31 |
| ctactgtgaaggtcctgaga | Dll1 exon_8 | atatgcctctcactgaggtc | Dll1 exon_32 |
| taagagtaccggaggtctgt | Dll1 exon_9 | agcagcagcaggaggacaag | Dll1 exon_33 |
| agtgctcgtcacacacaaac | Dll1 exon_10 | tggtgtttctgtagcttcag | Dll1 exon_34 |
| gaacacagagcaaccttctc | Dll1 exon_11 | ttgttcatggtttctgtctc | Dll1 exon_35 |
| cgcaggtgaagtggccaaag | Dll1 exon_12 | gctaacagaaacgtccttct | Dll1 exon_36 |
| gatgcactcatcgcagtagc | Dll1 exon_13 | ggtgttcttgatctgggtag | Dll1 exon_37 |
| catggagacaacctgggtat | Dll1 exon_14 | cgtgaaagtccgccttcttg | Dll1 exon_38 |
| cagttacactgccagggttg | Dll1 exon_15 | cttaaagctgctcttctcgg | Dll1 exon_39 |
| gtcggcaggaacatgtgtag | Dll1 exon_16 | atagtccacagtggggtatc | Dll1 exon_40 |
| cacagttggcacctgtatac | Dll1 exon_17 | ccttgaggtctcgaacgagg | Dll1 exon_41 |
| taggagcacactcatctact | Dll1 exon_18 | tcacgtttgctgtgtgtatc | Dll1 exon_42 |
| gagggcaggtgcaagagaag | Dll1 exon_19 | agctctgtgactggcacttg | Dll1 exon_43 |
| cacagaccttgccatagaag | Dll1 exon_20 | tttttctgtcaggaatctcc | Dll1 exon_44 |
| attgaagcaagggccatctg | Dll1 exon_21 | gtagagtagacagactctgg | Dll1 exon_45 |
| cagggttatctgaacatcgt | Dll1 exon_22 | ctggtacttggtgtcctttg | Dll1 exon_46 |
| tgaagccagagaagcccaag | Dll1 exon_23 | ctgcagacagaacatacacc | Dll1 exon_47 |
| gagatccatcttcttctcac | Dll1 exon_24 | tcgctataacacactcatcc | Dll1 exon_48 |

**Supplemental Table S7 - Dll1 intron smRNA FISH probe sequences**

| | | | |
|---|---|---|---|
| cccagtaagaagtcaagacc | Dll1 Intron_1 | gacgttggaaggaaatgcca | Dll1 Intron_25 |
| gacctttcaggtaagtagga | Dll1 Intron_2 | aaagtgactcgatatcctcc | Dll1 Intron_26 |
| ggggagagtacttgatggag | Dll1 Intron_3 | gcgctcaaaggatatgggaa | Dll1 Intron_27 |
| aactcctatcactttcctta | Dll1 Intron_4 | aaattcttggccaagacggg | Dll1 Intron_28 |
| aactcctatcactttcctta | Dll1 Intron_5 | cctgagagaggaaagctact | Dll1 Intron_29 |
| gagaaccaagtacagaggct | Dll1 Intron_6 | agtttagatttctagccacc | Dll1 Intron_30 |
| actgtgtatcaatggggaga | Dll1 Intron_7 | accccagtgtttgaataaga | Dll1 Intron_31 |
| gcagagtgaaggtaagaggc | Dll1 Intron_8 | tatggcacgtgtaagtaggg | Dll1 Intron_32 |
| agagagtggtatggatgctg | Dll1 Intron_9 | cctatcctcagagagatcta | Dll1 Intron_33 |
| gataacacctagacgctagg | Dll1 Intron_10 | gataagagtggacagctggt | Dll1 Intron_34 |
| agaagatgctcaggttggta | Dll1 Intron_11 | gtgcgtctaggacaaaaggg | Dll1 Intron_35 |
| acggccttagcaaaacactg | Dll1 Intron_12 | tggctttggacttcattcag | Dll1 Intron_36 |
| taaaatgccagaatccagcc | Dll1 Intron_13 | aaagctcctgagcaacgtag | Dll1 Intron_37 |
| cttcctacagtggacacaag | Dll1 Intron_14 | tactcaactaactccctgga | Dll1 Intron_38 |
| ctcctccctgtgaagaaaag | Dll1 Intron_15 | gccttcagtagagtaatagc | Dll1 Intron_39 |
| aatacaaaggccctagggag | Dll1 Intron_16 | acctatgctcgtacattttc | Dll1 Intron_40 |
| acattgttgcagcaaggaga | Dll1 Intron_17 | gctttttaactgaagcctgt | Dll1 Intron_41 |
| actctgtggaggaaagggag | Dll1 Intron_18 | agagtctcagatgaagggca | Dll1 Intron_42 |
| cccagggaatctttggaaac | Dll1 Intron_19 | caggacatcaggccatttaa | Dll1 Intron_43 |
| gaaggacagggcacacagag | Dll1 Intron_20 | cataagctacctgtccttaa | Dll1 Intron_44 |
| ccacaatcccaaggattgaa | Dll1 Intron_21 | caccaaaattatcctgcagc | Dll1 Intron_45 |
| taccctggtttgcaagaaga | Dll1 Intron_22 | gagacctggcaatcttagag | Dll1 Intron_46 |
| ctcagattgctggagagagg | Dll1 Intron_23 | caggaaggatctatgtttcc | Dll1 Intron_47 |
| ctacagattgggcagatgag | Dll1 Intron_24 | aggatcttcagtcagcacag | Dll1 Intron_48 |

**Supplemental Table S8 - Nestin exon smRNA FISH probe sequences**

| | | | |
|---|---|---|---|
| cagcgctgatcgacgaggac | Nestin exon_1 | ctccttgtcaagagatctga | Nestin exon_25 |
| gtctcaagggtattaggcaa | Nestin exon_2 | tccagtgattctatgttctc | Nestin exon_26 |
| tgtcttcagaaaggctgtca | Nestin exon_3 | tggctcttgagtatctacag | Nestin exon_27 |
| gttctggccttaaggaattc | Nestin exon_4 | ctattgtctcactagtcact | Nestin exon_28 |
| tctgacctctgcatttttag | Nestin exon_5 | ttgggtctcatcttctagag | Nestin exon_29 |
| agcagggaatgggggacatc | Nestin exon_6 | cttggttctcatccacacac | Nestin exon_30 |
| caaagaggctctggagcctg | Nestin exon_7 | ctttcaaggggtgtcagcac | Nestin exon_31 |
| gaggacaccagtagaactgg | Nestin exon_8 | agatctcagttcttgactct | Nestin exon_32 |
| gcctctaaaatagagtggtg | Nestin exon_9 | agagacctcagagactcttg | Nestin exon_33 |
| gaaactctagactcactgga | Nestin exon_10 | agatgcaactctgccttatc | Nestin exon_34 |
| ctttctttacaagttcccag | Nestin exon_11 | tgagcaactgggacctctag | Nestin exon_35 |
| ttctacttttacctctgtgg | Nestin exon_12 | ctctacggagtcttgttcac | Nestin exon_36 |
| cactttcttgtatttcctgt | Nestin exon_13 | aaatgccttgggtcctctag | Nestin exon_37 |
| gaatcctggatttcttctgt | Nestin exon_14 | tgatctatctttgccttcac | Nestin exon_38 |
| cagggtttccatctgcaaag | Nestin exon_15 | cttgcattccagagtctcac | Nestin exon_39 |
| agacatcagtggctcatctc | Nestin exon_16 | ctcatcttcctctgtcttag | Nestin exon_40 |
| tctcatggttctgggttttc | Nestin exon_17 | ttgggaccagggactgttag | Nestin exon_41 |
| ccgagttctcttctatagat | Nestin exon_18 | agatcgggatgggtgaacag | Nestin exon_42 |
| tgactctcgttctctagctt | Nestin exon_19 | tggctcatcttctactcttc | Nestin exon_43 |
| attctctttttccagagacc | Nestin exon_20 | gaatctccccacgaccaaac | Nestin exon_44 |
| ctagagacctcagggactct | Nestin exon_21 | aagggagtagagtcagggag | Nestin exon_45 |
| tcgagtttctgttcctcttg | Nestin exon_22 | caggagacttcaggtagagg | Nestin exon_46 |
| ttttctggagggttcactgt | Nestin exon_23 | cagatgagaggtcagagtca | Nestin exon_47 |
| ctttggtaactctggatcca | Nestin exon_24 | agatgatcggacacctcttt | Nestin exon_48 |

**Supplemental Table S9 - Nestin intron smRNA FISH probe sequences**

| | | | |
|---|---|---|---|
| cttggttttaccagggacag | Nestin Intron_1 | gcatagacactggagattgc | Nestin Intron_25 |
| ggactcctgaatacagttgt | Nestin Intron_2 | caaggtctctttcctgagac | Nestin Intron_26 |
| gacagggaagtctaggactt | Nestin Intron_3 | tggacacaaactacccacag | Nestin Intron_27 |
| tgctaggaggtctcagaatt | Nestin Intron_4 | cagtggattctggagaaggg | Nestin Intron_28 |
| agcagctacagtgggtattg | Nestin Intron_5 | tagatctgtctctggtcttg | Nestin Intron_29 |
| ggttgccattcaaagagtga | Nestin Intron_6 | attcagtccttcctagacac | Nestin Intron_30 |
| accttctgattgacaactgc | Nestin Intron_7 | gagcacatctggttctcaaa | Nestin Intron_31 |
| tggggagagagcatgtgttg | Nestin Intron_8 | tcagttcacaagggattcgg | Nestin Intron_32 |
| tatggttgctaggaaccttc | Nestin Intron_9 | agaggtctttaggttcacag | Nestin Intron_33 |
| gagaagccttgacacagaga | Nestin Intron_10 | gatcacaaatcctccttcag | Nestin Intron_34 |
| catctgtgcaatatgagggg | Nestin Intron_11 | gaagactgactggggcgaag | Nestin Intron_35 |
| caagaggctgttttctcagc | Nestin Intron_12 | gcaaacaccagtgcttacat | Nestin Intron_36 |
| caagaagagtgggcaggtgg | Nestin Intron_13 | aagtgggaattctcaggctg | Nestin Intron_37 |
| gagaggagatgggttacaca | Nestin Intron_14 | tttgtccacactaccatctg | Nestin Intron_38 |
| cacgggaggagatgggataa | Nestin Intron_15 | gatcagactcctcagatcag | Nestin Intron_39 |
| atgctattgagaacctagct | Nestin Intron_16 | ggccaagtgaattgctaaca | Nestin Intron_40 |
| tcttcttaggatgctggtta | Nestin Intron_17 | cagattcccacattcggaag | Nestin Intron_41 |
| caagcacaggaaacactcca | Nestin Intron_18 | ccatatttgagactcaggga | Nestin Intron_42 |
| cactgctcacatctttgtag | Nestin Intron_19 | cccacagatatatacttggt | Nestin Intron_43 |
| gatctctgggactcataagc | Nestin Intron_20 | tgtcgtatgaaagccacaca | Nestin Intron_44 |
| ctcaagtttatgcctaccaa | Nestin Intron_21 | gcaggatcactctttattgt | Nestin Intron_45 |
| ataaccatctcttctagtcc | Nestin Intron_22 | aagacttggactgatgctgc | Nestin Intron_46 |
| cctgaagactgagttgtctc | Nestin Intron_23 | actgcttagactggggacat | Nestin Intron_47 |
| agaactcgagagcaggtgac | Nestin Intron_24 | tcatttcagccattttatcc | Nestin Intron_48 |
